# A melon (*Cucumis melo*) homologue of REPRESSOR OF PHOTOSYNTHETIC GENES prevents chloroplast differentiation in the fruit flesh

**DOI:** 10.1101/2025.08.28.672820

**Authors:** Laura Valverde, Salvador Torres-Montilla, Marta Pujol, Jordi Garcia-Mas, Manuel Rodriguez-Concepcion

## Abstract

Fruit flesh color in melon can be orange, green or white, depending on the accumulation of the orange carotenoid β-carotene or / and green chlorophylls. The dominant allele of *Green flesh* (*Gf*) causes orange melons, but in the absence of this allele the flesh of ripe melon can be white or green depending on the *White flesh* (*Wf*) locus, being white dominant over green. The identity of *Wf* has remained unclear despite several candidates have been proposed. Here we identified *Wf* by fine mapping of a segregating population derived from the white-fleshed variety Piel de Sapo (PS, *gf gf / Wf Wf*) and the orange-fleshed Védrantais (VED, *Gf Gf / wf wf*). *Wf* corresponds to the gene *MELO3C003098*, herein referred to as *CmRPGE1* as it encodes a fruit-specific homologue of REPRESSOR OF PHOTOSYNTHETIC GENES (RPGE) microproteins. Similar to RPGE homologues from other plants, overexpression of the PS allele (*CmRPGE1*^*PS*^) caused a pale green leaf phenotype in *Nicotiana benthamiana* and *Arabidopsis thaliana*. By contrast, a 10-nucleotide deletion in the VED allele (*CmRPGE1*^*VED*^) resulted in a loss of RPGE function. The active CmRPGE1^PS^ microprotein interacts with a fruit-localized melon homologue of ARABIDOPSIS PSEUDO-RESPONSE REGULATOR2 (APRR2), a GARP family transcription factor. Binding of CmRPGE1^PS^ retains the melon APRR2 homologue in the cytosol, hence preventing the regulation of target genes involved in chloroplast biogenesis. In green fruit cultivars, the non-functional CmRPGE1^VED^ allele allows APRR2 to perform its function, leading to chloroplast development and consequently a green flesh phenotype.

**SIGNIFICANT STATEMENT:** Fruit color is a critical quality trait that directly influences consumer preference and market value in melon. In this study, we demonstrate that the *White flesh (Wf)* gene, a major determinant of melon fruit flesh color, encodes a fruit-specific homologue of REPRESSOR OF PHOTOSYNTHETIC GENES (RPGE) microproteins. The encoded microprotein, CmRPGE1, interacts with melon homologues of ARABIDOPSIS PSEUDO-RESPONSE REGULATOR2 (APRR2), a GARP family transcription factor. In white flesh melons, the CmAPRR2.2 paralog is sequestered in the cytoplasm upon interaction with CmRPGE1, thereby preventing the expression of target genes involved in chlorophyll accumulation. Conversely, the truncated version of CmRPGE1 present in green-fleshed melons fails to interact with CmAPRR2.2 and to prevent its nuclear localization and function, allowing chloroplasts differentiation.

## INTRODUCTION

The distinctive colors of ripe fleshy fruits attract animals to eat them and disperse the mature seeds in nature, but rind and flesh pigmentation is also an important quality trait for consumers. In melon (*Cucumis melo* L.), fruit flesh can be orange, green or white, depending on the accumulation of the orange carotenoid β-carotene and green chlorophylls. The loci *Green flesh* (*Gf*) and *White flesh* (*Wf*) are the main determinants of melon flesh color, being *Gf* epistatic over *Wf. Gf* is the major locus qualitatively differentiating between orange and non-orange flesh. It is located on chromosome 9 (Cuevas et al., 2009) and it corresponds to the gene *MELO3C005449*, also known as *ORANGE* (*CmOR*). A single amino acid substitution from Arg to His at position 108 of the CmOR protein induces the accumulation of β-carotene exclusively in the fruit, leading to the orange flesh coloration (Tzuri et al., 2015). Besides promoting the activity of phytoene synthase, the first and main-determining enzyme of the carotenoid pathway, the CmOR^His^ variant inhibits β-carotene hydroxylation and degradation by an unknown mechanism (Chayut et al., 2015; Tzuri et al., 2015; Chayut et al., 2017).

In the absence of the dominant *CmOr*^*His*^ allele, the flesh of ripe melon can be white or green depending on *Wf*, being white dominant over green. The *Wf* gene is located on chromosome 8 (Cuevas et al., 2009), but its identity remains unknown despite several candidates have been proposed. QTL mapping for fruit traits in a recombinant inbred line (RIL) population between PI 424723 (*C. melo* var. *momordica*, pale orange) and Dulce (*C. melo* var. *reticulatus*, deep orange) suggested *MELO3C003069* as a candidate for *Wf* (Galpaz et al., 2018). The encoded protein, CmPPR1 (PENTATRICOPEPTIDE REPEAT-CONTAINING PROTEIN 1), belongs to the PPR protein family, involved in plastid RNA processing. However, no fine mapping was conducted to define the region associated with this trait, nor was the gene functionally validated (Galpaz et al., 2018). Analysis of another RIL collection generated by crossing the white-fleshed melon variety Piel de Sapo (PS, *gf gf / Wf Wf*) and the orange-fleshed Védrantais (VED, *Gf Gf / wf wf*) allowed to identify a QTL on chromosome 8 associated with white and green flesh color, *LUMQU8.1* (*MELO3C003082* to *MELO3C003110*) (Pereira et al. 2021). Combining RIL genetic mapping with GWAS data narrowed the identified QTL interval to a region containing 11 annotated genes (*MELO3C003096* to *MELO3C003109*) but not *CmPPR1*, which was located 202 kb away (Zhao et al., 2019). It was proposed that one gene of this interval, *MELO3C003097*, could be a candidate for *Wf* as it encodes a homologue of SLOW GREEN 1 (SG1), a TETRATRICOPEPTIDE REPEAT-CONTAINING PROTEIN (TPR) family protein required for chloroplast development (Hu et al., 2014). Expression of this gene, referred to as *CmSG1*, was found to be higher in green-flesh compared to white-flesh accessions (Zhao et al., 2019). Similar to CmPPR1, however, it was unclear how a gain of function could result in a white flesh color and no further experiments were carried out to confirm the proposed role of CmSG1 in the pigmentation of melon flesh. Interestingly, the same peak on chromosome 8 was found in a GWAS analysis for both rind and flesh color (Zhao et al., 2019), suggesting an important role for the pigmentation of different melon fruit tissues. GWAS approaches further showed a pivotal role of the *MELO3C003375* gene in shaping fruit color variation in the rind but also in the flesh of melon fruit (Oren et al., 2019; Zhao et al., 2019). The encoded protein, CmAPRR2.1, is a homolog of the transcription factor APRR2 (ARABIDOPSIS PSEUDO-RESPONSE REGULATOR2-LIKE), a GARP (Golden2, ARR-B, Psr1) family member related to the GLK (GOLDEN2-LIKE) master activator of chloroplast development and chlorophyll biosynthesis (Chen et al., 2016). *APRR2* genes have been shown to regulate chlorophyll and carotenoid accumulation and green pigmentation in the fruits of many plant species, including cucurbits such as melon (Pan et al., 2013; Liu et al., 2016; Oren et al. 2019; Jeong et al., 2020; Ma et al., 2021; Arrones et al., 2022; Fang et al., 2023; Huo et al., 2023; Zhan et al., 2023; Cao et al. 2024; Gebretsadik et al., 2024; Guo et al.,2024; Wang et a., 2024; Chen et al., 2025; Fan et al., 2025). However, *MELO3C003375* cannot be *Wf* because it is located on chromosome 4 instead of chromosome 8.

Here we aimed at identifying the *Wf* gene through a fine mapping approach and characterizing the biological function of the encoded protein. We demonstrate that *Wf* is *MELO3C003098*, a gene encoding a homologue of REPRESSOR OF PHOTOSYNTHETIC GENES (RPGE) microproteins that prevents chlorophyll biosynthesis and chloroplast development in the fruit flesh by binding to APRR2 proteins to prevent their transcriptional activity.

## RESULTS

### Pigment accumulation and plastid differentiation differ in white and green melons

To facilitate the study of the genetic control of melon fruit flesh coloration, we used previously generated introgression lines (ILs) derived from the cross between the white-fleshed melon Piel de Sapo (PS, *gf gf / Wf Wf*) and the orange-fleshed Védrantais (VED, *Gf Gf / wf wf*) with different flesh colors (Pereira et al. 2021; Santo Domingo et al. 2022). In particular, we identified two ILs in the PS background showing VED genome introgressions in chromosome 8 with the recessive *wf* allele (VED8.3, *gf gf* / *wf wf*, green) or in chromosome 9 with the dominant *Gf* allele (VED9.2, *Gf Gf* / *Wf Wf*, orange), and two in the VED background showing PS genome introgressions in chromosome 8 with *Wf* (PS8.2, *Gf Gf* / *Wf Wf*, orange) and chromosome 9 with *gf* (PS9.3, *gf gf* / *wf wf*, green). The flesh color phenotype of these ILs (Figure 1 and Supplemental Figure S1) demonstrated that, unlike that proposed by some previous studies (Galpaz et al., 2018; Oren et al., 2019), the white and green flesh coloration trait is monogenic, and it also confirmed that *Gf* is epistatic over *Wf*.

**Figure 1.**
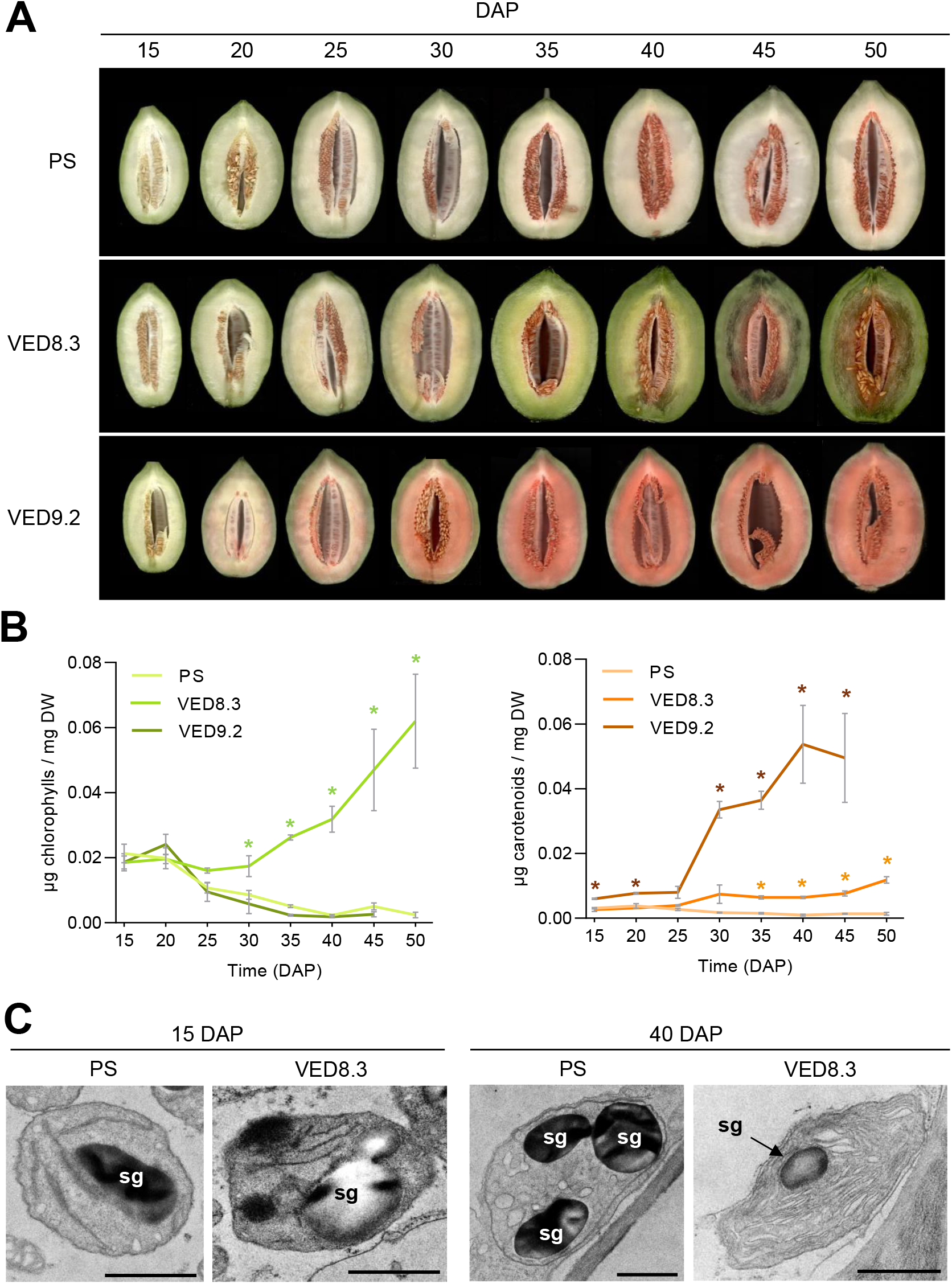
Melon flesh pigmentation and plastid ultrastructure during fruit development. (A) Longitudinal section of representative PS, VED8.3 and VED9.2 fruits collected at the indicated times (DAP, days after pollination). (B) Levels of chlorophylls and carotenoids in the flesh of the collected fruits. Mean and SEM values of three independent fruit (n=3) are shown. Asterisks mark statistically significant differences relative to PS samples at each time point according to t-test (P < 0.05). (C) Ultrastructure of melon flesh plastids from PS or VED8.3 fruits collected at 15 DAP (both pale green) and 40 DAP (white and green, respectively). Bar, 1 μm; sg, starch grain. All samples from 2020 harvest.

Next, we used the selected lines to monitor the development of flesh pigmentation during melon fruit ripening. Similar results were obtained in two different harvest years, 2020 (Figure 1 and Supplemental Figure S1) and 2021 (Supplemental Figure S2). Analysis of fruit development from 15 days after pollination (DAP) until harvest showed that all lines exhibited a pale green flesh coloration and low amounts of chlorophyll and carotenoid pigments during the early stages of development (15 DAP). Chlorophyll content decreased from 20 DAP to reach almost undetectable levels at 30 DAP in PS background lines with white (PS) or orange (VED9.2) flesh. In contrast, chlorophyll content remained constant up to 30 DAP in the green flesh VED8.3 line and then increased to reach the highest levels at harvest (50 DAP) (Figure 1A-B). A similar trend was observed in the VED background (Supplemental Figures S1 and S2). In the case of carotenoids, at 15 DAP they were already slightly increased in orange flesh lines VED9.2, VED and PS8.2 compared to white and green flesh lines of the same backgrounds (Figure 1A-B and Supplemental Figures S1 and S2). At later stages of fruit development, they decreased in the white flesh PS. While carotenoid contents increased in green flesh fruit VED8.3 and PS9.3, orange flesh fruit VED9.2, VED and PS8.2 showed a much more dramatic rise consistent with their characteristic flesh color (Figure 1A-B and Supplemental Figures S1 and S2).

We also analyzed plastid ultrastructure in the flesh of PS (white) and VED8.3 (green) lines at 15 and 40 DAP (Figure 1C). At 15 DAP, both lines showed plastids with large starch grains and poorly developed thylakoids. By contrast, different plastid structures were present in the flesh of PS and VED8.3 melons at 40 DAP, when the flesh color change was complete. PS plastids were similar to amyloplasts, as they contained large starch granules and vesicular structures instead of thylakoidal membranes (Figure 1C). In contrast, the plastids of ripe green VED8.3 melons exhibited a highly developed membrane system reminiscent of thylakoidal sacks but lacking grana (Figure 1C). Together, these results indicate that ripening involves the conversion of the plastids present in young fruit into amyloplasts in white-fleshed melons harboring the dominant *Wf* allele and rudimentary chloroplasts in green-fleshed (*wf wf*) melons.

### Fine mapping of *LUMQU8.1* singled out *MELO3C003098* as *Wf*

To identify the *Wf* gene, we performed a fine mapping in a BC1 population obtained by crossing PS (*Wf Wf*, white) with VED8.3 (*wf wf*, green) and subsequently backcrossing the generated F1 hybrid (*Wf wf*, white) with the VED8.3 parental line. QTL *LUMQU8.1* was originally mapped to a genetic interval of 3.6 cM, corresponding to a physical distance of 0.25 Mb that contains 32 annotated genes in the reference genome DHL92 v3.6.1 (Ruggieri et al., 2018; Pereira et al., 2018; Zhao et al., 2019). To narrow down the QTL region, a set of 26 SNPs was generated and tested in the parental lines (PS and VED8.3), the F1 hybrid and VED (Supplemental Figure S3). This analysis revealed that the VED introgression in VED8.3 finished between SNP markers *chr08:32019418* and *chr08:32023068*, reducing the original QTL interval to a region of 127 kb. The genotype of 2,500 BC1 plants allowed the identification of 40 recombinant individuals in this interval, which were then analyzed with the additional SNPs inside the interval (Supplemental Table S1).

For the phenotypic screening, we first analyzed the color of the ovary of the recombinant individuals, as we observed that PS and hybrid ovaries had a pale green color, while those of VED8.3 were darker green. We also analyzed the fruit flesh color at 40 DAP, confirming the association between ovary and fruit flesh color (i.e., pale green ovaries develop into fruits with white flesh, while dark green ovaries give rise to green flesh fruit) (Supplemental Figure S3). By associating the phenotype with the genotype, we were able to narrow down the QTL to 19.3 kb between markers *ch8:32003744* and *ch8: 32023068*. Within this region, there were four SNPs linked to the flesh coloration (*chr08:32010934, chr08:32012692, chr08:32013195* and *chr08:32019418*, which were targeting *MELO3C003098* (Supplemental Figure S3). These results conclusively demonstrated that the only complete gene in the initial *LUMQU8.1* interval associated with the white and green flesh coloration, and thus the candidate for *Wf*, is *MELO3C003098*.

### The PS allele of *MELO3C003098* encodes an active RPGE microprotein

*MELO3C003098* encodes a short protein (microprotein) with homology to a family of repressors of chloroplast development named REPRESSOR OF PHOTOSYNTHETIC GENES (RPGE) (Supplemental Figure S4). Besides *MELO3C003098* (here referred to as *CmRPGE1*), the melon genome contains three more RPGE homologues, namely *CmRPGE2* (*MELO3C025477*), *CmRPGE3* (*MELO3C023827*), and *CmRPGE4* (*MELO3C010673*) (Supplemental Figure S4A). Only *CmRPGE1* is fruit-specific (Supplemental Figure S4B). *CmRPGE2* and *CmRPGE3* are mainly expressed in leaves, although they also show expression in fruits. *CmRPGE4* is predominantly expressed in seedlings (Supplemental Figure S4B). Our RT-qPCRanalysis confirmed that *CmRPGE1* expression increased in the fruit flesh during the early stages of development in both PS and VED cultivars, but earlier induction and lower transcripts levels were detected in PS (Supplemental Figure S5).

Sequencing of the *CmRPGE1* gene in PS and VED genomes showed that the PS allele (from herein referred to as *CmRPGE1*^*PS*^) encodes a protein of 97 aa, whereas the VED allele (*CmRPGE1*^*VED*^) shows a 10 pb deletion in the region encoding the N-terminal domain of the protein, resulting in a predicted shorter protein of 86 aa (Supplemental Figure S6). Comparison to other CmRPGE paralogs from melon and homologs from other plants (Supplemental Figure S4A) suggested that CmRPGE1^PS^ likely corresponds to the full-length protein whereas CmRPGE1^VED^ might be a truncated variant. Overexpression of RPGE proteins has been shown to cause a pale phenotype in leaves (Ichikawa et al., 2006; Kim et al., 2016; Zhang et al., 2021; Kim et al., 2023; Han et al., 2024; Tachibana et al., 2024). We therefore tested the functionality of melon *CmRPGE1*^*PS*^ and *CmRPGE1*^*VED*^ alleles by transiently expressing them in *Nicotiana benthamiana* leaves (Figure 2). Three days after agroinfiltration with constructs harboring the genomic *CmRPGE1* sequence from PS or VED either alone or fused to the GFP marker protein (Figure 2A), only the areas expressing the CmRPGE1^PS^ variants acquired a pale color (Figure 2B) and showed reduced levels of photosynthetic pigments (Figure 2C) without triggering a senescence process (Figure 2D). By contrast, leaf areas expressing the CmRPGE1^VED^ variants were undistinguishable from controls agroinfiltrated with an empty vector or the GFP marker alone. These results support the conclusion that only the CmRPGE1^PS^ variant is biologically active.

**Figure 2.**
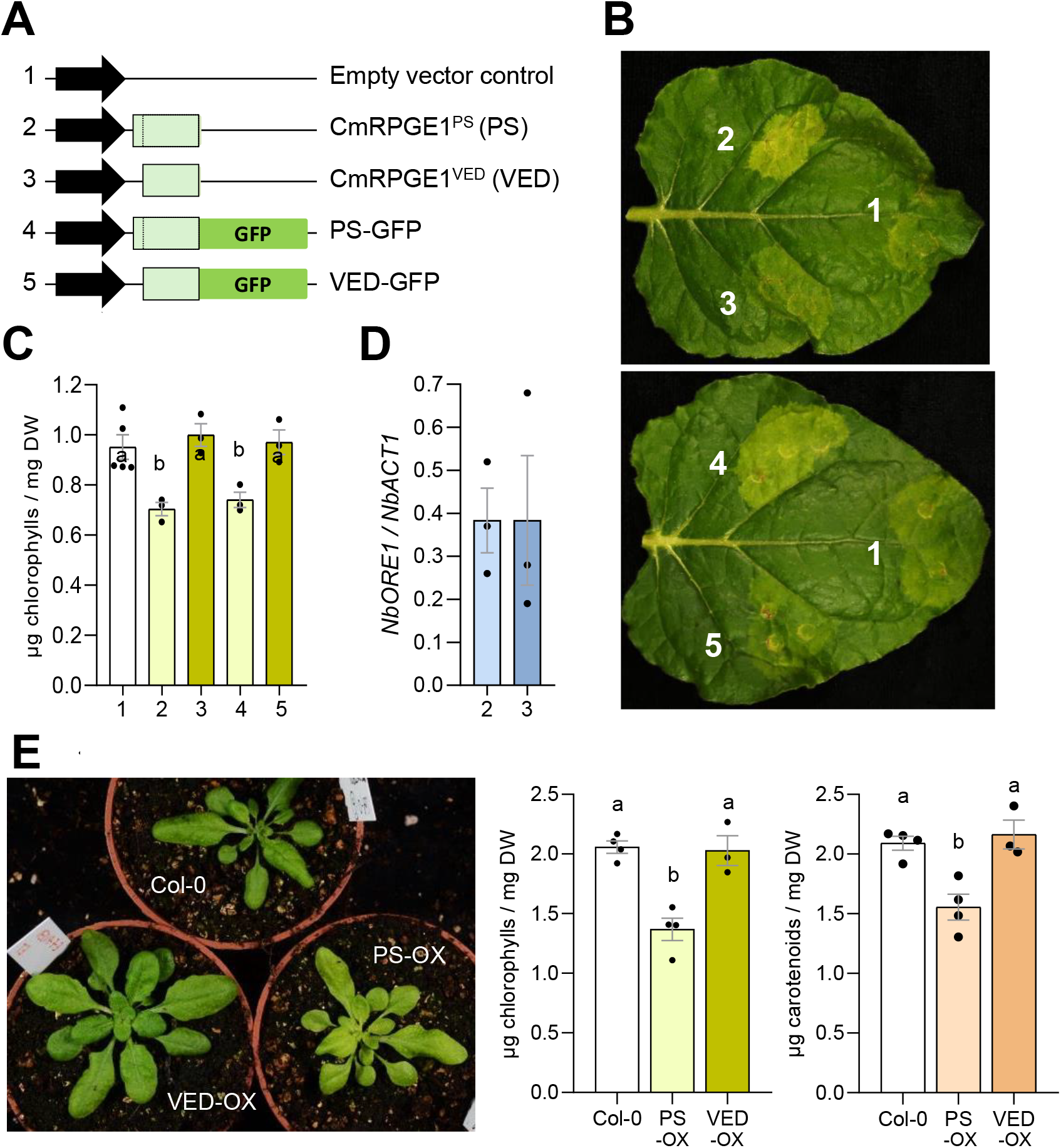
Only the PS variant of CmRPGE1 cause a pale leaf phenotype. (A) Constructs used for agroinfiltration of *N. benthamiana* leaves. Melon sequences correspond to genomic DNA. (B) Images of representative leaves three days after agroinfiltration of the indicated leaf sections with different constructs. (C) Chlorophyll content of the agroinfiltrated leaf sections. Mean and SEM of three sections from different leaves (n=3) are shown. (D) Levels of *N. benthamiana ORE1* transcripts in leaves agroinfiltrated with the indicated constructs. Data correspond to the mean and SEM of RT-qPCR analysis of three independent samples (n=3). (E) Representative images of Arabidopsis plants constitutively overexpressing genomic copies of *CmRPGE1*^*PS*^ (PS-OX) or *CmRPGE1*^*VED*^ (VED-OX) and the untransformed wild-type (Col-0). Plot on the right shows the mean and SEM of photosynthetic pigment contents in the rosette leaves of three independent plants (n=3). Letters in (C) and (D) represent statistic significances determined by one-way ANOVA followed by Tukey’s multiple comparison test, P < 0.05).

Confocal analysis of the leaf areas expressing the GFP-tagged proteins showed a similar accumulation of CmRPGE1^PS^-GFP and CmRPGE1^VED^-GFP proteins in the cytosol and the nucleus of mesophyll cells (Supplemental Figure S7), similar to that reported for other RPGE homologs (Zhang et al., 2021; Tachibana et al., 2024). Co-infiltration of CmRPGE1^VED^-GFP together with a RFP-tagged version of CmRPGE1^PS^ showed full overlapping of GFP and RFP fluorescence (Supplemental Figure S7), indicating that both PS and VED variants localize similarly in the cell despite their differences in the N-terminal region and derived impact on functionality.

### CmRPGE1^PS^ functions similar to the Arabidopsis RPGE proteins

To further validate the functional role of the CmRPGE proteins, the genomic sequences of the PS and VED variants were stably expressed in Arabidopsis. Homozygous lines confirmed to overexpress *CmRPGE1*^*PS*^ (named as PS-OX) or *CmRPGE1*^*VED*^ (VED-OX) were selected for experiments. Consistent with the results of the transient expression experiments (Figure 2A-D), a pale phenotype associated to decreased levels of photosynthetic pigments (chlorophylls and carotenoids) was observed in PS-OX but not in VED-OX lines (Figure 2E and Supplemental Figure S8). To confirm that this pale phenotype did not result from a senescence-related degradation of chloroplasts, we carried out dark-induced senescence experiments using plate-grown seedlings of the transgenic lines grown together with untransformed wild-type (Col-0) controls. No differences in visual phenotype or photosynthetic pigment loss were observed among the three genotypes following incubation of up to 21 days in darkness (Supplemental Figure S9). By contrast, deetiolation experiments involving the light-triggered differentiation of chloroplasts from the etioplasts present in dark-grown seedlings showed a similar greening for Col-0 and VED-OX seedlings but a substantially reduced rate in the case of PS-OX plants (Supplemental Figure S10A). As a consequence, the levels of chlorophylls and carotenoids 24h after illumination of etiolated seedlings were lower in PS-OX than in VED-OX or Col-0 (Supplemental Figure S10A). RNA-seq analysis of the same samples confirmed that overexpression of *CmRPGE1*^*PS*^ (but not *CmRPGE1*^*VED*^) caused a substantial decrease in the level of transcripts for components of the photosynthetic apparatus and for enzymes involved in the production of photosynthetic pigments or in the Calvin cycle (Supplemental Figure S10B). These results demonstrate that the pale phenotype associated to CmRPGE1^PS^ activity is not due to chloroplast degradation but results from inhibition of chloroplast differentiation.

Similar to melon, RPGE is also encoded by a small gene family in Arabidopsis (Kim et al., 2023). Four genes encode Arabidopsis RPGE-like proteins, namely AtRPGE1 (AT5G02580), AtRPGE2 (AT3G55240), AtRPGE3 (AT3G28990) and AtRPGE4 (AT1G10657). CmRPGE1^PS^ is most similar to AtRPGE2, also known as PEL (PSEUDO-ETIOLATION IN LIGHT) and BPG4 (BRZ-INSENSITIVE-PALE GREEN 4) due to the characteristic pale phenotype of overexpressing plants (Ichikawa et al., 2006; Tachibana et al., 2024). Overexpression of AtRPGE1 also results in a pale phenotype (Kim et al., 2016; Kim et al., 2023). Interestingly, almost half of the genes differentially expressed in light-grown Arabidopsis seedlings overexpressing AtRPGE2 compared to untransformed controls (Kim et al., 2023) were similarly misregulated in PS-OX seedlings exposed to light for 24h (Supplemental Figure S11). Many of the genes differentially expressed in both AtRPGE2-OX and PS-OX seedlings are related to photosynthesis and response to light, strongly suggesting that melon and Arabidopsis homologs share at least part of the molecular mechanism eventually causing the pale phenotype characteristic of overexpressing lines.

### Binding of CmRPGE1^PS^ to CmAPRR2.2 prevents its nuclear localization

RPGE proteins exert their function by binding to GARP family transcription factors such as GLK and APRR2, reportedly preventing their binding to DNA and the expression of target genes (Kim et al., 2023; Tachibana et al., 2024; Wang et al. 2024). We speculated that CmRPGE1^PS^ (but not CmRPGE1^VED^) might cause the white flesh phenotype by preventing the function of melon fruit GLK or/and APRR2 proteins. The melon genome contains a single gene encoding GLK (*MELO3C019337*), but it is not expressed in fruits (Supplemental Figure S12A) and its loss of function causes a general rather than a fruit-specific bleaching (Yang et al. 2023). In the case of APRR2, two genes encode paralogs with expression in the fruit flesh, referred to as CmAPRR2.1 (*MELO3C003375*) and CmAPRR2.2 (*MELO3C013874*) (Cao et al. 2024) (Supplemental Figure S12B). Our RT-qPCR analysis confirmed that both genes are expressed in the flesh during development of white PS and green VED8.3 fruit (Supplemental Figure S12C). At the protein sequence level, CmAPRR2.1 and CmAPRR2.2 are 54% identical. CmAPRR2.1 has been found to be associated to the green pigmentation of the melon fruit rind and suggested to also have a similar role in the flesh (Oren et al. 2019; Cao et al., 2024), whereas no information associating CmAPRR2.2 with the coloration of melon fruit tissues was available. Strikingly, CmAPRR2.2 is phylogenetically closer to homologs from Arabidopsis and carrot but also to other APRR2 proteins associated with the green pigmentation of fruit tissues in many plant species, including tomato, pepper, and cucurbits such as bitter gourd, cucumber, eggplant, pumpkin, zucchini and watermelon (Pan et al., 2013; Liu et al., 2016; Jeong et al., 2020; Fang et al., 2023; Gebretsadik et al., 2024; Guo et al., 2024; Wang et al. 2024; Chen et al., 2025; Fan et al., 2025) (Supplemental Figure S13).

The possible interaction of CmRPGE1 variants with CmAPRR2.1 and CmAPRR2.2 was initially tested by bimolecular fluorescence complementation (BiFC) in *N. benthamiana* leaves agroinfiltrated with the corresponding constructs (Figure 3). APRR2 paralogs were fused to the N-terminal half of YFP (NY-A2.1 and NY-A2.2) and CmRPGE1 variants were fused to the C-terminal half of this fluorescent protein (CY-PS and CY-VED) (Figure 3A). Reconstituted YFP fluorescence was observed in the nucleus of leaf cells co-expressing NY-A2.1 with either CY-PS or CY-VED (Figure 3B). By contrast, both nuclear and cytosolic fluorescence was detected with the NY-A2.2 and CY-PS combination and no signal could be obtained with when NY-A2.2 was co-expressed with CY-VED (Figure 3B). In the absence of CmRPGE1 proteins, RFP-tagged CmAPRR2.1 (A2.1-RFP) showed an almost exclusive localization in nuclei of agroinfiltrated *N. benthamiana* leaf cells, whereas a higher proportion of A2.2-RFP was evident in the cytosol (Figure 3C). The cytosolic localization of A2.2-RFP was further promoted in cells co-expressing a GFP-tagged CmRPGE^PS^ protein (PS-GFP) (Figure 3D), resulting in a RFP fluorescence distribution similar to the YFP fluorescence emitted upon interaction of NY-A2.2 and CY-PS (Figure 3B). In agreement with the absence of BiFC-detected interaction between CmAPRR2.2 and CmRPGE^VED^ (Figure 3B), the cellular distribution of A2.2-RFP fluorescence (i.e., abundant in nuclei but also present in the cytosol) remained unchanged in the presence of VED-GFP (Figure 3D).

**Figure 3.**
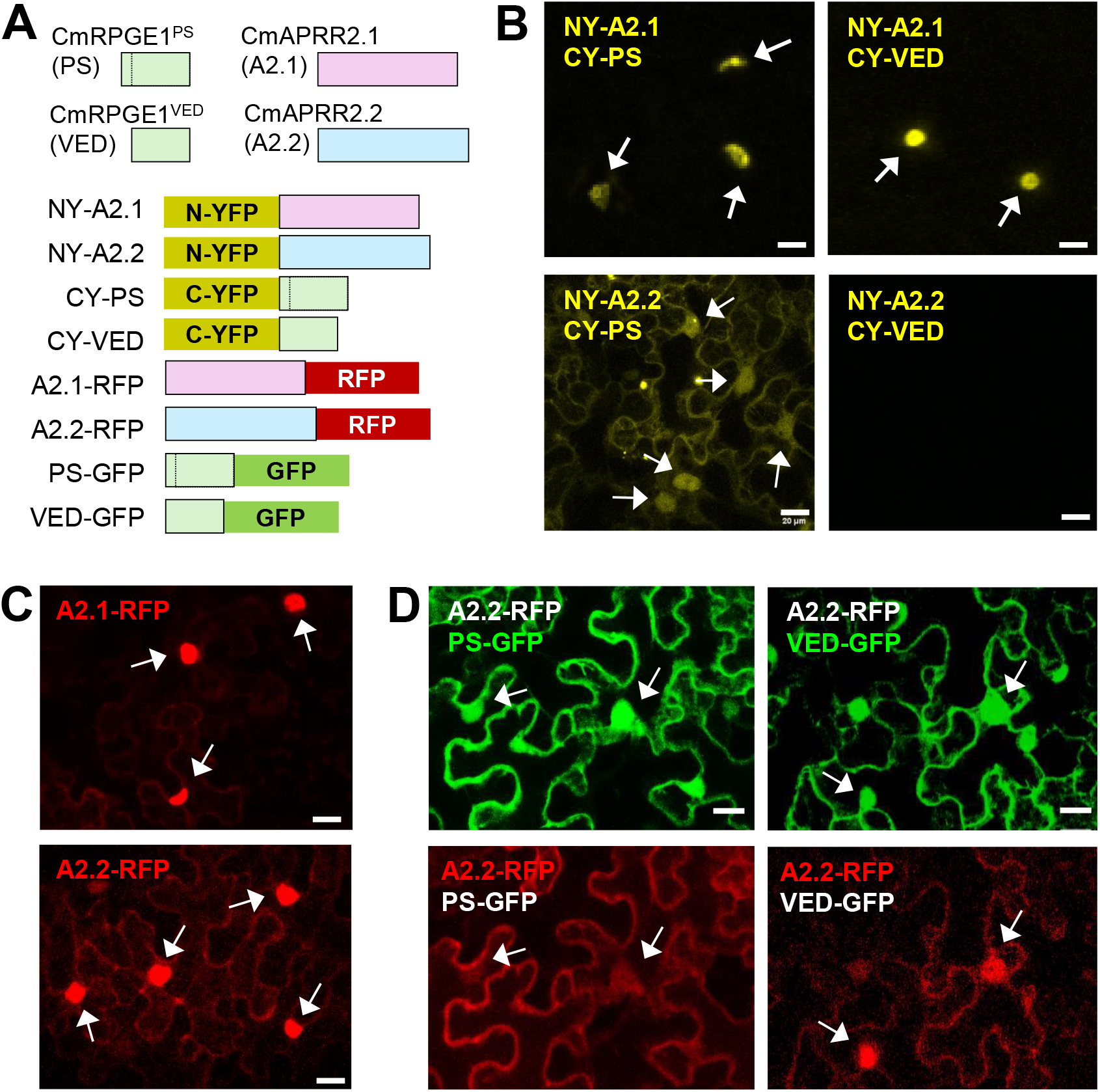
CmRPGE1 variants interact with CmAPRR2 paralogs in different cell compartments. (A) Constructs used for agroinfiltration of *N. benthamiana* leaves. Melon sequences correspond to cDNA. (B) Bimolecular Fluorescence Complementation (BiFC) assay with the indicated constructs. Yellow fluorescence indicates successful interaction. (C) Localization of RFP-fused CmAPRR2 proteins (in red). (D) Localization of GFP-fused CmRPGE1 variants (in green, upper panels) and RFP-fused CmAPRR2.2 proteins (in red, lower panels) when co-expressed in the same cells. Upper and lower panels of the same combination correspond to the same field. All confocal images were taken three days after agroinfiltration with the indicated constructs. Arrows mark the position of nuclei. Bars = 20 μm.

The results were next validated by co-immunoprecipitation experiments in *N. benthamiana* leaves agroinfiltrated with the corresponding constructs encoding the melon APRR2 paralogs with a C-terminal hemagglutinin tag (A2.1-HA and A2.2-HA) and the CmRPGE1 variants with a C-terminal myc tag (PS-myc and VED-myc) (Figure 4). Consistent with the BiFC results, A2.1-HA was immunoprecipited with both PS-myc and VED-myc whereas A2.2-HA could only be immunoprecipited with PS-myc (Figure 4A). Strikingly, A2.2-HA protein levels in co-agroinfiltrated leaves appeared to increase in the presence of PS-myc compared to VED-myc (Figure 4A). RT-qPCR analysis of the same agroinfiltrated leaves showed that *A2.2-HA* transcript levels were similar in VED-myc + A2.2-HA and PS-myc + A2.2-HA samples (Figure 4B), indicating that the CmRPGE1^PS^-mediated overaccumulation of CmAPRR2.2 protein is not due to increased gene expression. Additional experiments showed that co-expression of VED-myc had no effect on A2.2-HA protein levels compared to controls containing only A2.2-HA constructs, and confirmed that the presence of PS-myc led to higher A2.2-HA protein levels (Figure 4C). Together, the described results confirmed that CmRPGE1^PS^ binds to both CmAPRR2.1 and CmAPRR2.2 whereas the shorter CmRPGE1^VED^ variant is only able to interact with CmAPRR2.1. However, only CmRPGE1^PS^ binding to CmAPRR2.2 interferes with nuclear localization of this transcription factor, causing its accumulation in the cytosol presumably to repress the regulation of its target genes.

**Figure 4.**
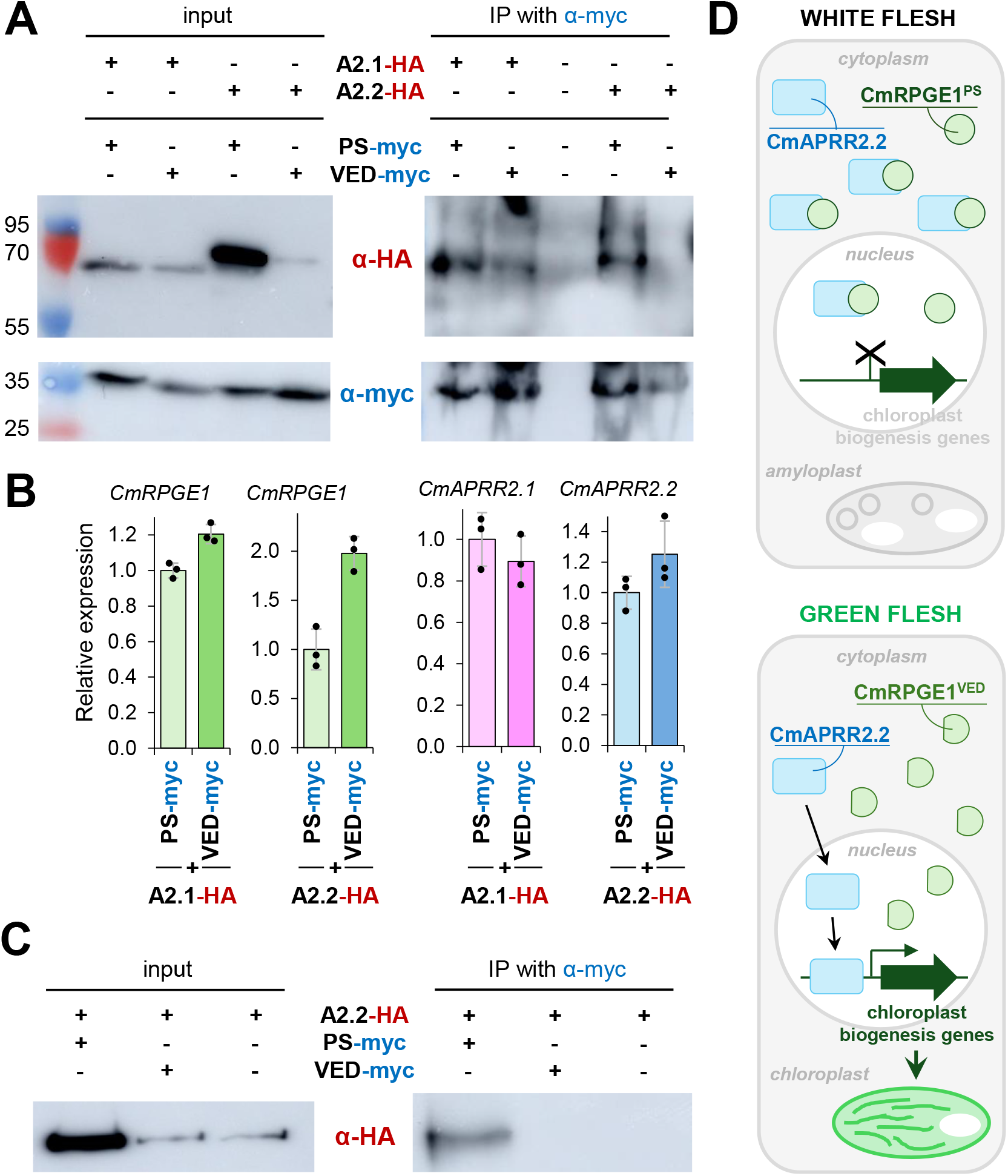
CmRPGE1^PS^ binding stabilizes CmAPRR2.2 protein. *N. benthamiana* leaves were co-agroinfiltrated with constructs encoding the PS and VED variants of CmRPGE1 with a C-terminal myc tag (PS-myc and VED-myc) and CmAPRR2 paralogs with C-terminal hemagglutinin (HA) epitopes (A2.1-HA and A2.2-HA). (A) Immunoblot analysis with antibodies against myc (blue) and HA (red) of protein extracts before (input) and after immunoprecipitation (IP) with anti-myc antibodies. Molecular weight (kDa) of protein standards is indicated in the left. (B) Transcript levels in the agroinfiltrated leaf samples. Data correspond to mean and SD values of RT-qPCR analysis of three independent samples (n=3), and they are represented relative to the levels in samples containing PS-myc. (C) Immunoblot analysis of a different experiment including samples without co-expressed CmRPGE1 variants. (D) Schematic model of the molecular control of flesh color in white and green melons. White flesh melons have a functional CmRPGEPS microprotein that binds to the transcription factor CmAPRR2.2 and sequesters it in the cytosol, hence preventing expression of target genes required for chloroplast differentiation. The shorter CmRPGEVED variant present in green flesh melons is unable to bind CmAPRR2.2, allowing its translocation to the nucleus to regulate the expression of chloroplast biogenesis genes.

## DISCUSSION

Despite the melon *Wf* gene was known for a long time to be located in chromosome 8, the efforts to identify its molecular function remained unsuccessful until now. Suggested candidate genes such as *MELO3C003069 / CmPPR1* (Galpaz et al., 2018) and *MELO3C003097 / CmSG1* (Zhao et al., 2019) were potentially required for chloroplast development. However, how the mutations detected in these genes could lead to the dominant nature of the *Wf* gene and the white color of the melon fruit flesh remained unclear and no experimental validation was provided. Additionally, GWAS approaches led to identify the melon *MELO3C003375 / CmAPRR2.1* gene as a major player in determining the green color of the fruit rind but also possibly the flesh (Oren et al., 2019; Zhao et al., 2019). CmAPRR2.1 shows homology to APRR2 transcription factors that positively regulate chloroplast development and green pigmentation of fruit tissues in many plant species (Pan et al., 2013; Liu et al., 2016; Jeong et al., 2020; Ma et al., 2021; Arrones et al., 2022; Fang et al., 2023; Huo et al., 2023; Gebretsadik et al., 2024; Guo et al., 2024; Chen et al., 2025; Fan et al., 2025). Strikingly, this gene is located in chromosome 4 instead of 8, and hence it was ruled out as *Wf*. Our work reported here reconciles these apparently contradictory observations by demonstrating that *Wf* is indeed a chromosome 8 gene, *MELO3C003098*, whose product (CmRPGE1) eventually regulates chlorophyll accumulation and chloroplast differentiation in the flesh by directly interfering with APRR2 function.

The function reported for Arabidopsis RPGE/PEL/BPG4 microproteins as negative regulators of chloroplast development (Ichikawa et al. 2006; Kim et al., 2016; Kim et al. 2023; Han et al. 2023; Tachibana et al. 2024) is conserved in rice (Zhang et al. 2021). Furthermore, comparison of transcriptomes of seedlings overexpressing RPGE proteins from Arabidopsis and melon strongly suggested a conserved mode of action of the functional CmRPGE^PS^ variant (Supplemental Figure S11). Work on the mode of action of RPGE has shown that these microproteins repress photosynthetic development by binding to GLK transcription factors, preventing the formation of active dimers (Kim et al. 2023) and their binding to DNA (Kim et al. 2023; Tachibana et al. 2024). Binding of the carrot RPGE homologue DcRPGE1, encoded by the *Y* locus (*DCAR_032551*), to the GLK-related protein DcAPRR2 (*DCAR_008297*) was also found to impair DcAPRR2 binding to its target DNA sequences (Wang et al. 2024). In all the cases, BiFC assays showed that RPGE-GLK and RPGE-APRR2 interactions predominantly occurred in the nucleus, even though some fluorescence signal could also be detected in the cytosol (Zhang et al. 2021; Kim et al. 2023; Tachibana et al. 2024; Wang et al. 2024). By contrast, our BiFC results show that the melon CmPRGE1^PS^ variant, shown to be functionally active based on its ability to cause a pale phenotype when overexpressed (Figures 3 and 4), interacts with the CmAPRR2.2 protein in both cytosolic and nuclear locations (Figure 3B). Most strikingly, this interaction promotes the cytosolic retention of CmAPRR2.2, hence preventing its nuclear activity as a transcription factor (Figure 3). This mechanism expands the functional scope of the RPGE microproteins beyond its previously reported role as nuclear suppressors of DNA binding of GARP family transcription factors to include cytoplasmic sequestration of these transcription factors. Strikingly, this mechanism does not apply for CmAPRR2.1, which unlike CmAPRR2.2 is almost exclusively localized in nuclei, where it is able to interact with both active CmPRGE1^PS^ and non-functional CmPRGE1^VED^ (Figure 3). Another distinctive feature of CmAPRR2.2 is that protein levels are highly increased upon interaction with CmPRGE1^PS^ (Figure 4). How cytosolic retention eventually results in enhanced CmAPRR2.2 levels remains unknown but we speculate that these transcription factors might be steadily degraded in the nucleus under physiological conditions to prevent a too long, prolonged response.

Interestingly, the same peak on chromosome 8 was found in GWAS for both rind and flesh color (Zhao et al., 2019). It is likely that this peak corresponds to the *CmRPGE1* gene, suggesting an important role for this microprotein in the pigmentation of different melon fruit tissues. A possible contribution of CmRPGE1 to melon rind coloration would likely involve interaction with CmAPRR2.1, which has been demonstrated to determine green rind pigmentation in melon (Oren et al. 2019; Cao et al. 2024; Chen et al., 2025). However, the observation that CmAPRR2.1 binds to functional (PS) and non-functional (VED) variants of the CmRPGE1 protein and that CmAPRR2.1 protein levels or localization are independent of the presence of the microproteins (Figures 5 and 6) strongly suggests that CmRPGE1 proteins might not regulate CmAPRR2.1 activity *in vivo*. Further work would be required to address this question.

Based on the experimental evidence available from our work and elsewhere, we propose a molecular regulation model of fruit flesh color in melon (Figure 4D). During the first stages of development, melon fruits are pale green and have plastids with poorly differentiated membranous structures (Figure 1 and Supplemental Figure S1). It is likely that these membranous structures are the sites harboring the low amounts of chlorophylls detected in these fruits (Figure 1 and Supplemental Figure S1). In orange-fleshed melons, the activity of the *Gf* gene (*CmOr*^*His*^ variant) induces carotenoid synthesis and chromoplast differentiation (Tzuri et al. 2015), hence preventing the formation of chloroplasts or amyloplasts. In lines lacking this variant, ripe fruits will be white or green depending on the *CmRPGE1* variant. Despite the active *CmRPGE*^*PS*^ variant present in white flesh melons and the non-functional *CmRPGE*^*VED*^ variant of green cultivars show different expression levels, they both peak at 30 DAP (Supplemental Figure S5), paralleling the first color changes associated with amyloplast (white) or chloroplast (green) differentiation (Figure 1 and Supplemental Figure S1). We propose that, at this stage, CmRPGE1^PS^ interacts with the CmAPRR2.2 protein and retains it in the cytoplasm of flesh cells, preventing the regulation of its target genes (Figures 5-7). While the specific physiological role of CmAPRR2.2 has not been demonstrated, we can deduce a role in promoting chlorophyll synthesis and chloroplast development based on its similarity with other APRR2 homologs associated with the green pigmentation of fruits from a variety of plant species of the families Solanaceae (Pan et al., 2013; Jeong et al., 2020; Arrones et al., 2022; Fang et al., 2023) and Cucurbitaceae (Liu et al., 2016; Ma et al., 2021; Zhan et al., 2023; Huo et al., 2023; Gebretsadik et al., 2024; Guo et al.,2024; Fan et al., 2025), including melon (Oren et al. 2019; Cao et al. 2024; Chen et al., 2025). In green fruit cultivars, the non-functional CmRPGE1^VED^ protein allows CmAPRR2.2 to perform its function, leading to a green flesh phenotype (Figure 4D).

Fruit color is a key trait that not only influences consumer preference and market value but also reflects complex biochemical processes, including those involved in the synthesis of health-promoting compounds such as carotenoids. Understanding the basic molecular mechanisms that regulate fruit color, particularly in melons, is also crucial for addressing a central question in biology, i.e., how plastids differentiate into specialized types such as chloroplasts, amyloplasts, and chromoplasts. Elucidating the genetic and molecular pathways responsible for these processes represents a critical step toward improving agricultural practices and crop quality, including the development of biofortified fruits with greater nutritional benefits.

## MATERIALS AND METHODS

### Plant material and growth conditions

Melon (*Cucumis melo*) ‘Piel de Sapo’ T111 (PS), ‘Védrantais’ (VED) and introgression lines PS8.2, PS9.3, VED8.3 and VED9.2 were already available in the lab (Pereira et al. 2021; Santo Domingo et al. 2022). Seeds were germinated and grown under greenhouse conditions in Torre Marimon (Caldes de Montbui, Barcelona) as described (Santo Domingo et al. 2024). Fruit collection for flesh color analysis was carried out from April to August in two consecutive years, 2020 and 2021. Samples were taken from the plant every 5 days from 15 DAP (Days After Pollination) until harvest (35-40 DAP in VED background and 50-55 in PS background). Fruits were cut open longitudinally and flesh images were taken with a scanner For fine mapping, parental lines PS and VED8.3 were crossed and a F1 hybrid was grown from August to November 2020 to be backcrossed with the parental VED8.3. Approximately 2500 plants of the backcross population (BC1) generated from March to June 2021 were genotyped for recombinant selection and afterwards grown in the greenhouse. Ovary coloration was analyzed during flowering. All plants were self-pollinated and fruit flesh color was determined at 40-45 DAP (August-November 2021). *Nicotiana benthamiana* plants used for transient expression experiments were grown in the greenhouse under long day conditions (LD, 14h of light at 26±1ºC and 10h of darkness at 21±1ºC). *Arabidopsis thaliana* Columbia (Col-0) ecotype was used to generate transgenic lines. For dark-induced senescence assays, Col-0 and transgenic lines were germinated and grown for 14 days at 24ºC under a photoperiod of 16h of light (140 μmol·m^-2^·s^-1^) and 8h of darkness. Then, the plates with the seedlings were placed in continuous darkness by wrapping them in aluminium foil. For deetiolation assays, seeds were stratified for 4 days in the dark and then illuminated for 2h to promote simultaneous germination. Then, plates were wrapped in aluminium foil and incubated at 24ºC for 3 days. Immediately after removing the aluminium foil, plates with etiolated seedlings were placed in a growth chamber with continuous light (85 μmol·m^-2^·s^-1^) and samples were taken at different time points following illumination.

### DNA extraction and genotyping

Two different DNA extraction protocols were used, one for quick genotyping with SNPs (Lu et al. 2020) and another one for lor long-term storage and PCR genotyping (Pereira et al. 2020). Plants used for the fine mapping were genotyped using SNP markers by the PACE2.0 SNP genotyping system (3CR Bioscience, Harlow, UK) based on an allele-specific PCR with different fluorochromes depending on the allele. Primers were designed following manufacturer instructions and are listed in Supplementary Table S1. These primers were tested with the parental lines (PS and VED8.3) as well as with VED and the F1 hybrid as controls and then used for the fine mapping assay.

### Constructs

Primers used for PCR amplification and cloning are listed in Supplementary Table S2 and constructs are described in Supplementary Table S3. Genomic DNA (gDNA) was used to amplify the *CmRPGE1* coding sequence from PS and VED plants using specific primers with or without the translation stop codon. For some experiments, cDNA was used instead of gDNA to amplify target sequences encoding CmRPGE1^PS^ and CmRPGE1^VED^ but also CmAPRR2.1 and CmAPRR2.2. The resulting amplicons were cloned into pDONR207 entry plasmids using Gateway (GW) technology (Invitrogen). Following sequencing to confirm absence of undesired mutations, the gDNA sequences were subcloned through LR reactions into pGWB605 (with GFP) or pGWB454 (with RFP) for transient expression in *N. benthamiana* (Supplementary Table S3). The pGWB605 clones with the endogenous translation stop codon (i.e., encoding the CmRPGE1 variants not fused to GFP) were used for stable expression in *A. thaliana*. For co-immunoprecipitation assays, cDNA sequences were cloned into plasmids pGWB420 (with a myc tag) and pGWB414 (with a HA tag) (Supplementary Table S3). For Bimolecular Fluorescence Complementation assays, cDNA sequences were cloned into plasmids YFN43-GW and YFC43-GW (Belda-Palazon et al. 2012).

### Transient and stable expression

Transient expression was carried out by agroinfiltration of *N. benthamiana* leaves from 4-5 weeks old plants with the indicated constructs as described (Andersen et al. 2021). To prevent silencing, *N. benthamiana* leaves were co-infiltrated with *Agrobacterium tumefaciens* GV3101 cultures transformed with the appropriate constructs together with a second strain harboring a HcPro silencing suppressor. Cultures were mixed in identical proportions when agroinfiltrating construct combinations. Stable Arabidopsis lines were generated by floral dip as described (Clough and Bent 1998). Positive transformants were selected in MS media supplemented with 20 mg/mL BASTA. Four independent lines showing a single T-DNA insertion (as determined by the Mendelian segregation of BASTA resistance) were selected for each genotype and taken to homozygosis for further characterization.

### Chlorophyll and carotenoid measurements

Quantification of photosynthetic pigments in melon fruit flesh samples was done by spectrophotometry. Samples were extracted using 50 mg of freeze-dried tissue and 1 ml hexane:acetone:methanol (2:1:1) as extraction solvent. Samples were resuspended in 800 μl of acetone. Absorbance was measured at 470 nm, 644.8 nm and 661.6 nm and the amount of pigments was estimated using the following equations: Chlorophylls = 7.05 x [A661.6] -18.09 x [A644.8]; Chlorophyll *a* = 11.24 x [A661.6] – 2.04 x [A644.8]; Chlorophyll *b* = 20.13 x [A644.8] – 4.19 x [A661.6]; Carotenoids = (1000 x [A470] – 1.9 x [Chlorophyll *a*] – 63.14 x [Chlorophyll *b*]) / 214. Chlorophylls and carotenoids in leaf samples were extracted and quantified by HPLC-DAD as described (Barja et al. 2021).

### Microscopy

Plastid ultrastructure was analyzed using Transmission Electron Microscopy (TEM) of melon fruit flesh samples fixed for 2 h at room temperature in 100 mM sodium cacodylate buffer with 2.5 % glutaraldehyde and 2 % paraformaldehyde. After overnight incubation at 4°C and several washes with cacodylate buffer, samples were fixed in 1 % OsO_4_, cleared with water, dehydrated in an acetone gradient, and embedded in Spurr resin. After microtomy, ultrathin sections were stained with 2 % uranyl acetate followed by 2.6 % lead citrate. Samples were observed using a Tecnai F20 microscope, and images were recorded using Digital Micrograph 3.3.0 software. Subcellular localization of proteins fused to fluorescent proteins (GFP, RFP, YFP) was visualized three days after agroinfiltration of *N. benthamiana* leaves with the corresponding constructs. A Leica TCS SP8-MP confocal laser-scanning microscope was used to detect GFP and YFP fluorescence (BP55-525 filter after excitation at 488 nm) and RFP (laser excitement at 532 nm and detection at 588 nm). All images were acquired using the same confocal parameters.

### Immunoprecipitation assays

Two days after agroinfiltration with the corresponding constructs, leaf samples (about 2 g) were collected and frozen in liquid nitrogen. Protein extraction and immunoprecipitation were performed as described (Barja et al. 2021). The presence of Myc- and HA-tagged proteins in input and immunoprecipitated samples was detected by immunoblot analyses using a 1:10,000 dilution of αMyc-HRP (Invitrogen) and a 1:2,500 dilution of αHA-HRP (Roche). SuperSignal^TM^ West Pico PLUS Chemiluminiscent Substrate (Thermo Scientific) was used for detection and the signal was visualized using the ChemiDoc Touch Imaging System (Bio-Rad).

### RNA isolation and RT-qPCR analysis

Total RNA was extracted using Spectrum™ Plant Total RNA Kit (Sigma-Aldrich). DNAse treatment was done with the TURBO DNA-free Kit (Thermo Fisher Scientific). Retro-transcription to cDNA was performed using PrimeScript™ RT-PCR (Takara Bio INC), and qPCR experiments were carried out using LightCycler480 (Roche) and QuantStudio^TM^3 (Thermo Fisher Scientific) with SYBR Green MasterMix. Reference genes for normalization were melon *CmRPS15* (*MELO3C006471*) and *N. benthamiana NbACT1* (*AY594294*). Primers used for RT-qPCR are listed in Supplementary Table S2.

### RNA-seq

Total RNA was extracted from transgenic *A. thaliana* lines PS-OX and VED-OX germinated and grown in the dark for 3 days and then exposed to light for 24h. RNA sequencing, read processing, and basic data analysis were performed by Novogene. After quality control, samples were sequenced using the Illumina Novaseq platform using the PE150 (Paired-Ends) strategy. More than 6.235 million high-quality reads were obtained from each sample after processing and cleaning the raw reads. High-quality reads were mapped and assembled with the *A. thaliana* reference genome using Hisat2 v2.0.5 and StringTie (v1.3.3b) (Pertea et al. 2015). Approximately 95% of the reads were successfully aligned. Transcript levels were calculated using FPKM values. Differentially expressed genes (DEGs) were determined using the R package DESeq2 (1.20.0). Only genes with P values lower than 0.05 were considered differentially expressed. Heatmaps of specific genes were generated from FPKM values using the R package pheatmap. Analysis of enriched Gene Ontology (GO) terms was performed using the R package clusterProfiler.

### Bioinformatic tools and statistical analyses

Expression profiles of melon genes in different tissues of the plant and the fruit were extracted from Melonet-DB (https://melonet-db.dna.affrc.go.jp/ap/mvw), and they correspond to the green flesh melon cultivar ‘Harukei-3’. Search for APRR2 homologs was carried out in NCBI (https://www.ncbi.nlm.nih.gov/protein/) and PLAZA dicots 5.0 (https://bioinformatics.psb.ugent.be/plaza.dev/), and nucleotide sequences were translated into protein sequences using Reverse Complement (https://www.bioinformatics.org/sms/rev_comp.html). Alignments were done in Uniprot (https://www.uniprot.org/align) using the default parameters. Phylogenetic trees were constructed using ITOL (Interactive Tree Of Life, https://itol.embl.de/). Statistical analyses were performed using the packages available in GraphPad (https://www.graphpad.com/).

## ACKNOWLEDGEMENTS

We thank the Cryomicroscopy Unit of the Barcelona Science Park (Centres Científics i Tecnològics de la Universitat de Barcelona, CCITUB) for sample fixation and processing for TEM and M. Rosa Rodriguez-Goberna (CRAG Metabolomics Facility) for excellent technical support with HPLC. We also thank Jaume F. Martinez-Garcia for the BiFC vectors and Elena del Blanco for help with melon protocols.

