## Supplemental data for "A melon (*Cucumis melo*) homologue of REPRESSOR OF PHOTOSYNTHETIC GENES prevents chloroplast differentiation in the fruit flesh"

**SUPPLEMENTARY FIGURES**

**Figure S1. Pigment content of melon flesh during fruit development.** (A) Longitudinal section of representative VED, PS8.2 and PS9.3 fruits collected at the indicated times (DAP, days after pollination). (B) Levels of chlorophylls and carotenoids in the flesh of the collected fruits. Mean and SEM values of three independent fruit (n=3) are shown. Asterisks mark statistically significant differences relative to VED samples at each time point according to t-test (P < 0.05). Samples from harvest of 2020.

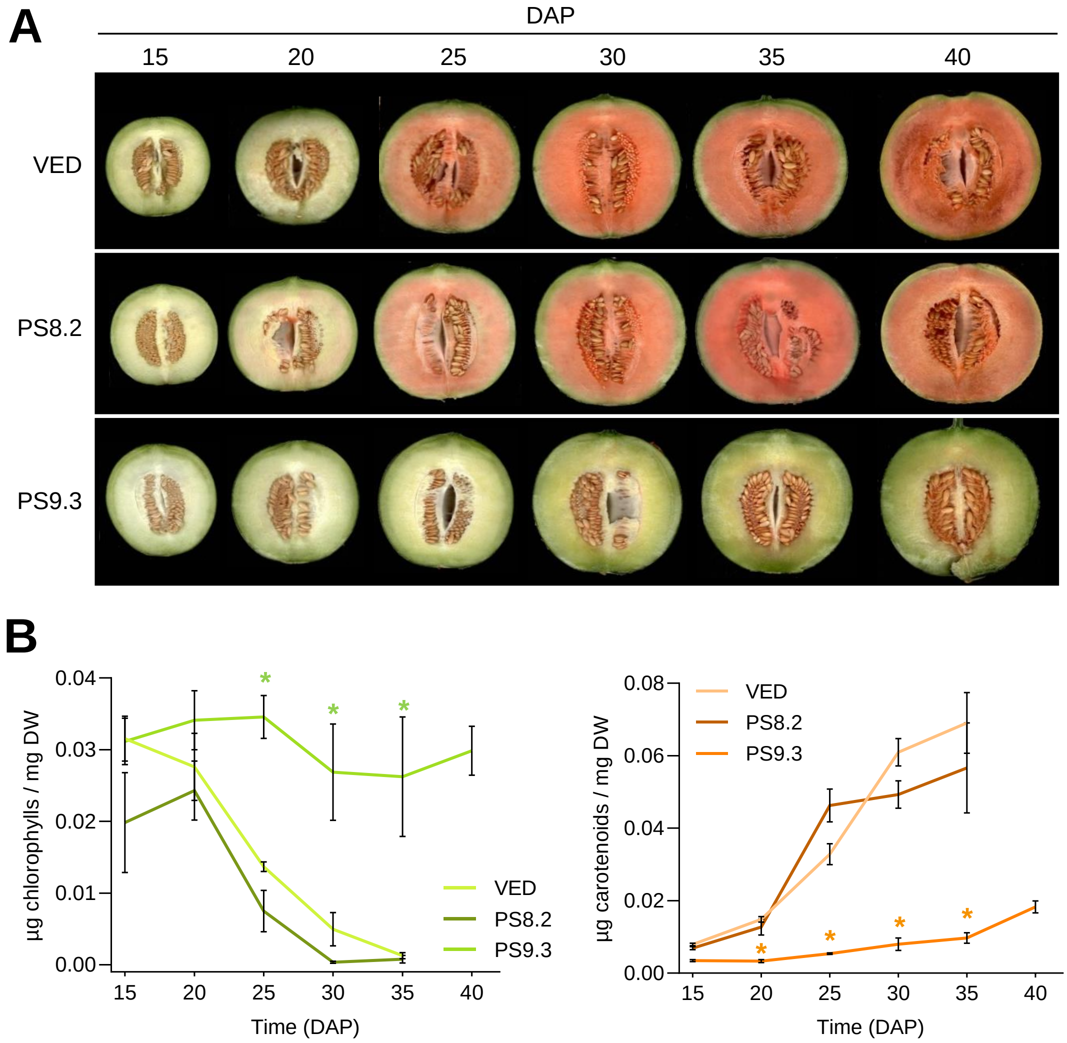

**Figure S2. Levels of chlorophylls and carotenoids in the flesh of the collected fruits of the indicated genotypes.** Mean and SEM values of three independent fruit (n=3) are shown. Asterisks mark statistically significant differences relative to PS (A) or VED (B) samples at each time point according to t-test (P < 0.05). Samples from harvest of 2021.

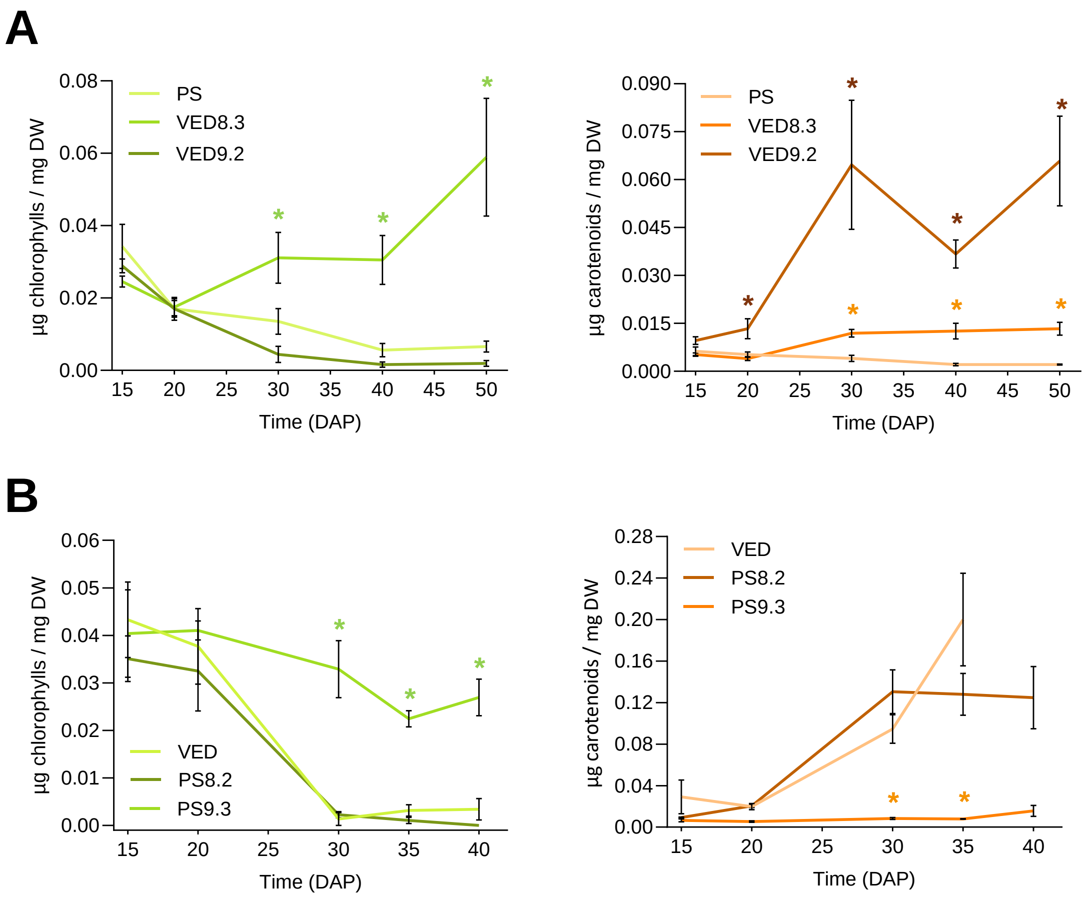

**Figure S3. Fine mapping of *LUMQU8.1* in the BC1 population.** Upper table represents the genotype of PS (white), VED (green), VED8.3 (green), and the hybrid (PS x VED8.3) (white) for the markers generated in the 0.25Mb region of the QTL *LUMQU8.1* (Melonomics v3.6.1). Grey, homozygous PS; Green, homozygous VED; Yellow, heterozygous. Lower Table represents the genotype-phenotype association assay of the 40 recombinant BC1 plants found in the region that comprises between markers chr08:31987002 and chr08:32019418, where *Wf* is located.

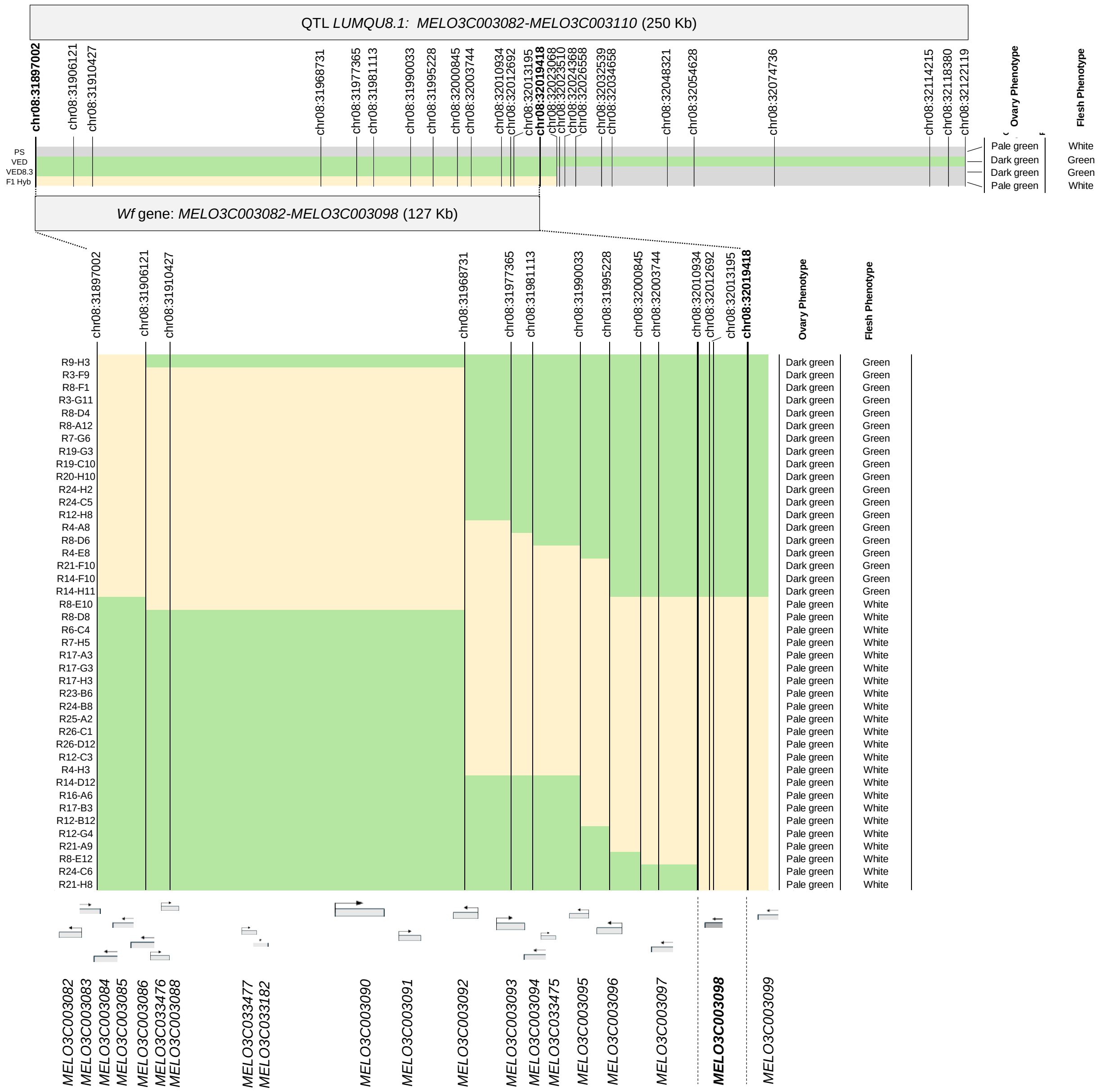

**Figure S4. The RPGE family in melon.** (A) Alignment of RPGE paralogs identified in melon with homologs with demonstrated roles in *Arabidopsis thaliana* (*AtRPGE1*, *AtRPGE2*), *Daucus carota* (*DcRPGE*, *DCAR_032551*) and *Oryza sativa* (*OsRPGE / DGP1*, *Os01g62060*). Alignment was performed using default parameters in Uniprot. Conserved residues are highlighted in blue. The conserved TIGR01589 domain is underlined. (B) Histogram representing expression levels of the four genes encoding RPGE homologs in different organs of the Harukei-3 melon variety (Melonet DB Gene Expression Atlas). Values in leaves and fruit flesh are boxed in green and black, respectively.

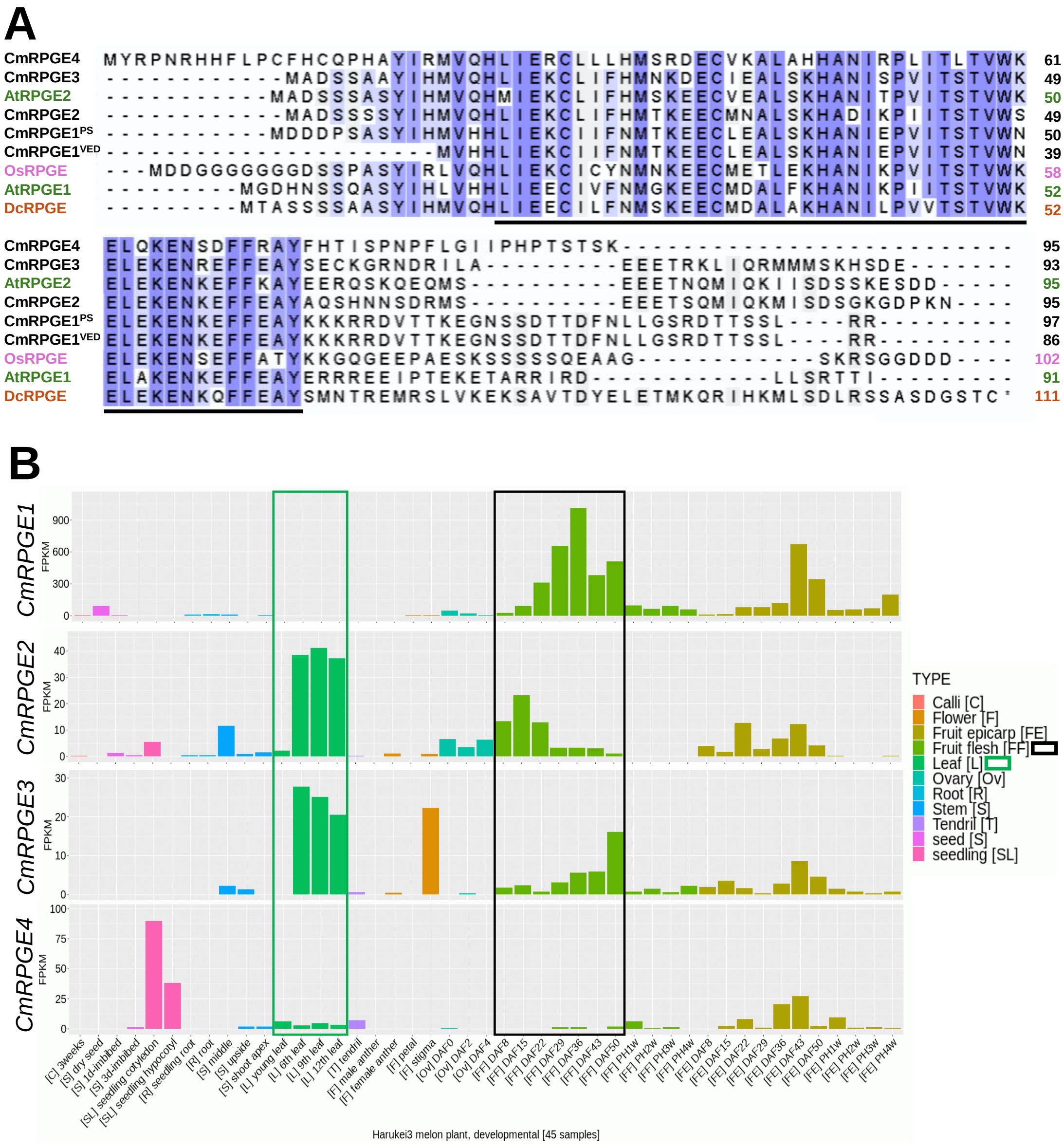

**Figure S5. Levels of *CmRPGE1* transcripts in the melon flesh during fruit development.** Data correspond to RT-qPCR analysis of samples from white (PS) and green (VED8.3) melons. Mean and SD values of three independent fruit samples (n=3) are shown. Asterisks mark statistically significant differences at each time point according to t-test (P < 0.01).

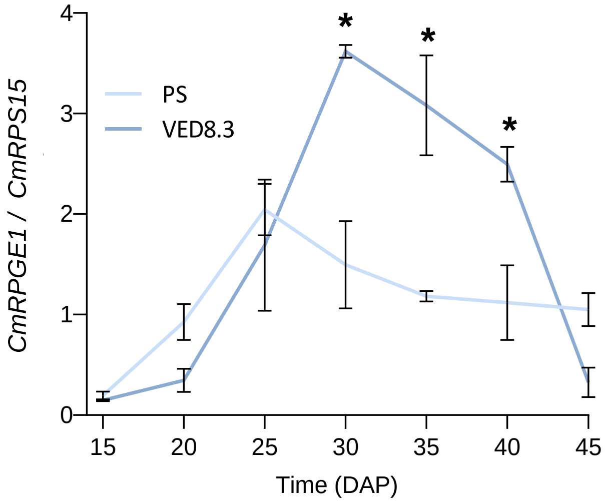


**Figure S6. DNA and protein sequences of the CmRPGE1 variants found in PS and VED melons.**

***CmRPGE1$^{PS}$***

```
acc atg gac gat gac ccc tct gct tca tac atc cac atg gtg cac cac ctg atc gaa aag
     M   D   D   D   P   S   A   S   Y   I   H   M   V   H   H   L   I   E   K
tgt ata att ttc aac atg act aaa gaa gag tgc ttg gaa gct ctc tcc aaa cat gca aac
 C   I   I   F   N   M   T   K   E   E   C   L   E   A   L   S   K   H   A   N
atc gaa ccg gtc atc act tcc acc gtg tgg aat gaa ttg gag aag gag aac aag gaa ttc
 I   E   P   V   I   T   S   T   V   W   N   E   L   E   K   E   N   K   E   F
ttc gaa gcg tat aag aag aaa cga cgt gac gtt act acg aag gag ggg aat tcg tcc gac
 F   E   A   Y   K   K   K   R   R   D   V   T   T   K   E   G   N   S   S   D
acg acg gat ttc aat ctg ctg ggt tcg agg gat acg acg tcg tct ctg agg cgg tag
 T   T   D   F   N   L   L   G   S   R   D   T   T   S   S   L   R   R   -
```

***CmRPGE1$^{VED}$***

```
Acc atg gac gat gac ccc t-- --- --- --c atc cac atg gtg cac cac ctg atc gaa aag
                                                 M   V   H   H   L   I   E   K
tgt ata att ttc aac atg act aaa gaa gag tgc ttg gaa gct ctc tcc aaa cat gca aac
 C   I   I   F   N   M   T   K   E   E   C   L   E   A   L   S   K   H   A   N
atc gaa ccg gtc atc act tcc acc gtg tgg aat gaa ttg gag aag gag aac aag gaa ttc
 I   E   P   V   I   T   S   T   V   W   N   E   L   E   K   E   N   K   E   F
ttc gaa gcg tat aag aag aaa cga cgt gac gtt act acg aag gag ggg aat tcg tcc gac
 F   E   A   Y   K   K   K   R   R   D   V   T   T   K   E   G   N   S   S   D
acg acg gat ttc aat ctg ctg ggt tcg agg gat acg acg tcg tct ctg agg cgg tag
 T   T   D   F   N   L   L   G   S   R   D   T   T   S   S   L   R   R   -
```

**Figure S7. The PS and VED variants of CmRPGE1 co-localize in the nucleus and the cytoplasm of plant cells.** Confocal microscopy pictures show the localization of the indicated fusion proteins three days after agroinfiltration of *N. benthaliama* leaves with the corresponding constructs. Green, GFP fluorescence; Red, RFP fluorescence; Yellow, overlapping GFP and RFP fluorescence.

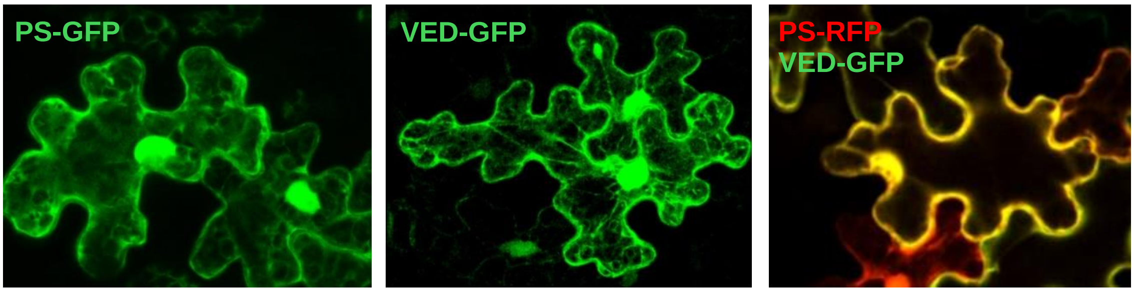


**Figure S8. Photosynthetic pigment contents in the rosette leaves of Col-0, PS-OX and VED-OX lines.** Mean and SEM of at least three independent homozygous individuals per line are shown. Letters represent statistic significances determined by one-way ANOVA followed by Tukey's multiple comparison test, P < 0.05).

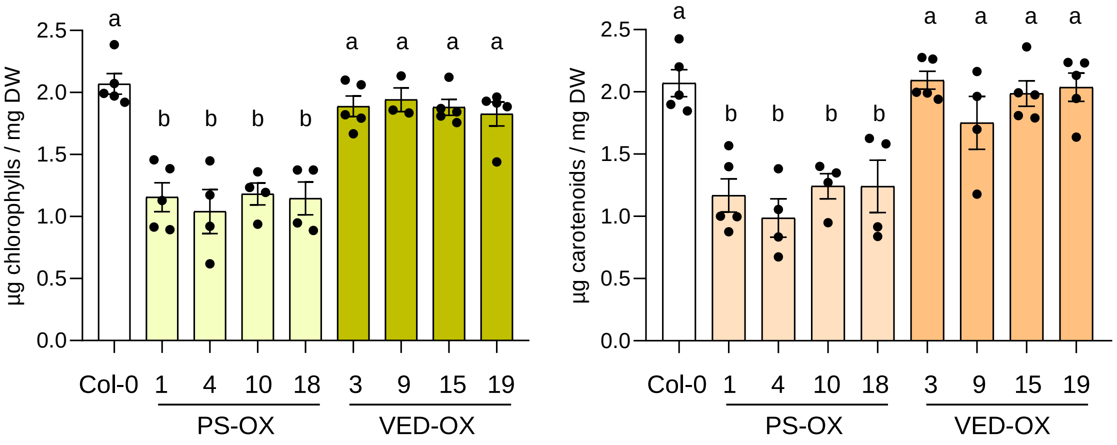

**Figure S9. Response of Col-0, PS-OX and VED-OX lines to dark-induced senescence.** (A) Representative images of seedlings of the indicated genotypes germinated and grown for two weeks in a long-day photoperiod and then incubated in the dark for 0, 7, 14 or 21 days. (B) Photosynthetic pigment contents of the seedlings treated as described in (A). Mean and SEM of three independent replicates (n=3) are shown. Data are shown relative to the pigment levels at the start of the experiment (D0).

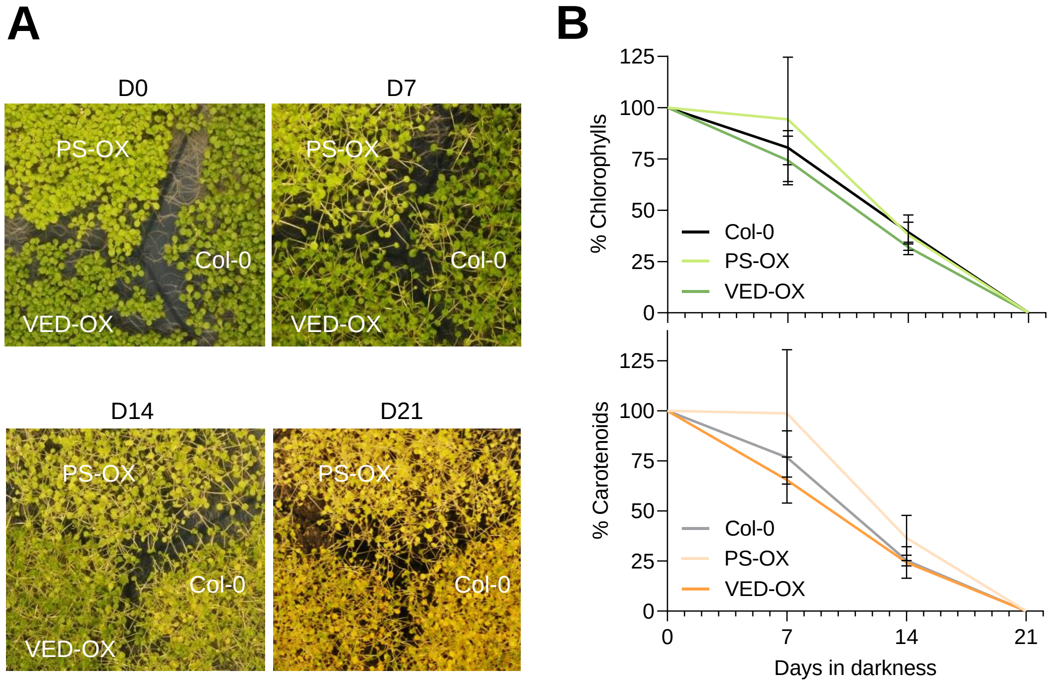

**Figure S10. CmRPGE1$^{PS}$ but not CmRPGE1$^{VED}$ prevents proper photosynthetic development in transgenic Arabidopsis plants.** (A) Photosynthetic pigment accumulation during deetiolation of the indicated lines. Data correspond to mean and SEM of four replicates (n=4). Asterisks mark statistically significant differences relative to Col-0 samples at each time point according to t-test ($P < 0.05$). (B) Heatmaps of the expression levels of genes involved in photosynthesis and photosynthetic pigment metabolism after 24h of deetiolation.

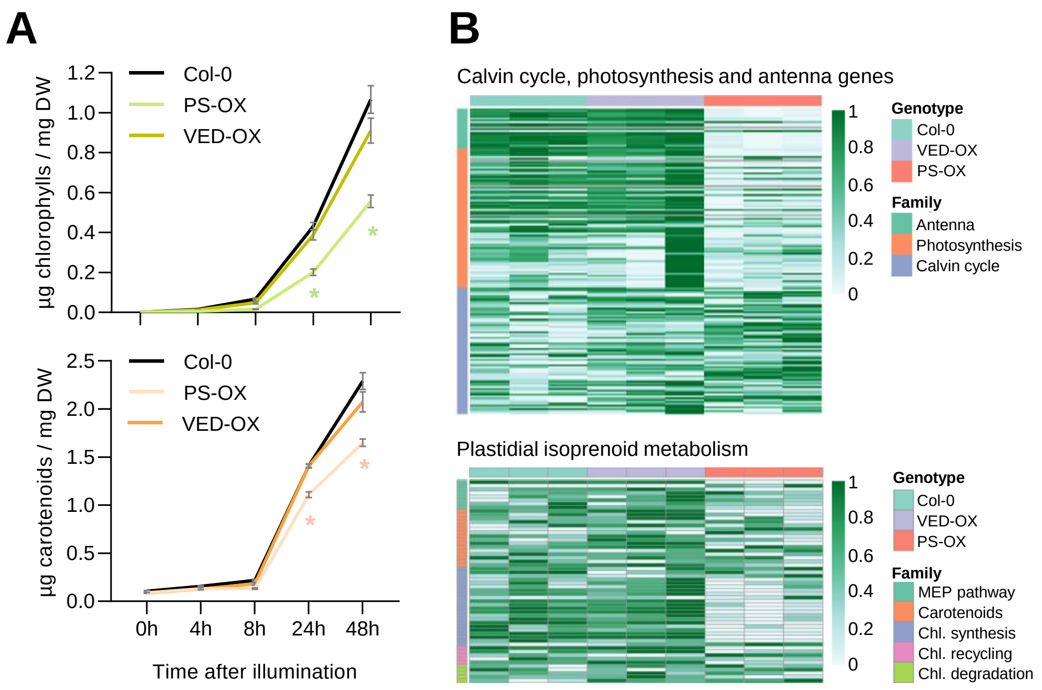

**Figure S11. AtRPGE2 and CmRPGE1$^{PS}$ have overlapping target genes.** (A) Venn diagram comparing differentially expressed genes (DEGs) in the RNAseq analyses of AtRPGE2-OX vs Col-0 and PS-OX vs Col-0. (B) Co-regulation of the 421 overlapping DEGs from the comparisons indicated in (A). (C) Gene ontology (GO) analysis of the 421 overlapping DEGs. The 10 most enriched Biological Processes are shown according to P value. The abundance of these DEGs in the indicated categories is compared to the proportion of Arabidopsis genes in the same categories.

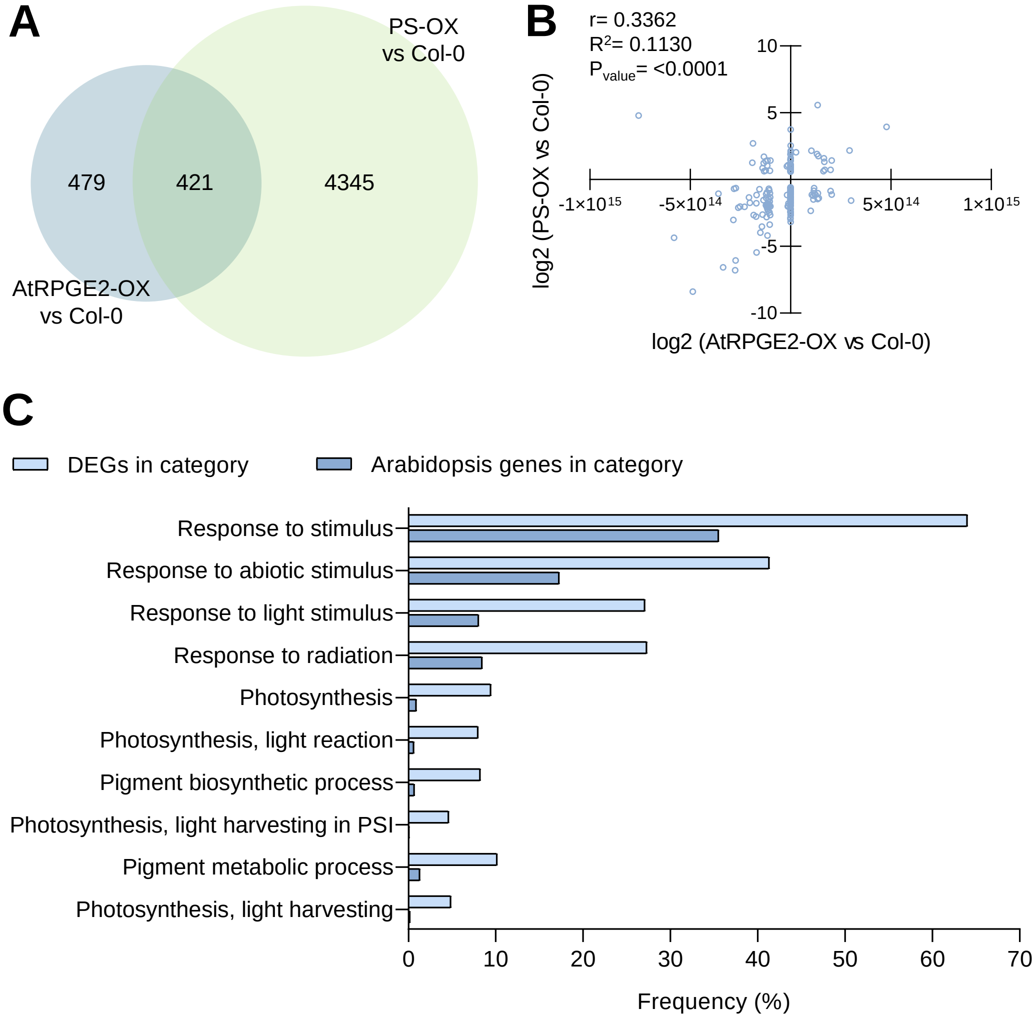

**Figure S12. Expression profiles of melon *CmGLK, CmAPRR2.1* and *CmAPRR2.2*.** (A) Distribution of *CmGLK* transcripts in different organs of the Harukei-3 melon variety according to the Melonet DB Gene Expression Atlas. (B) Distribution of *CmAPRR2.1* and *CmAPRR2.2* transcripts in different organs of Harukei-3 plants (Melonet DB Gene Expression Atlas). Histogram values in leaves and ripening fruit flesh are boxed in green and black, respectively. (C) Levels of *CmAPRR2.1* and *CmAPRR2.2* transcripts in the melon flesh during fruit development. Data correspond to RT-qPCR analysis of samples from white (PS) and green (VED8.3) melons. Mean and SD values of n=3 independent fruit samples are shown. Asterisks mark statistically significant differences at each time point according to t-test ($P < 0.01$).

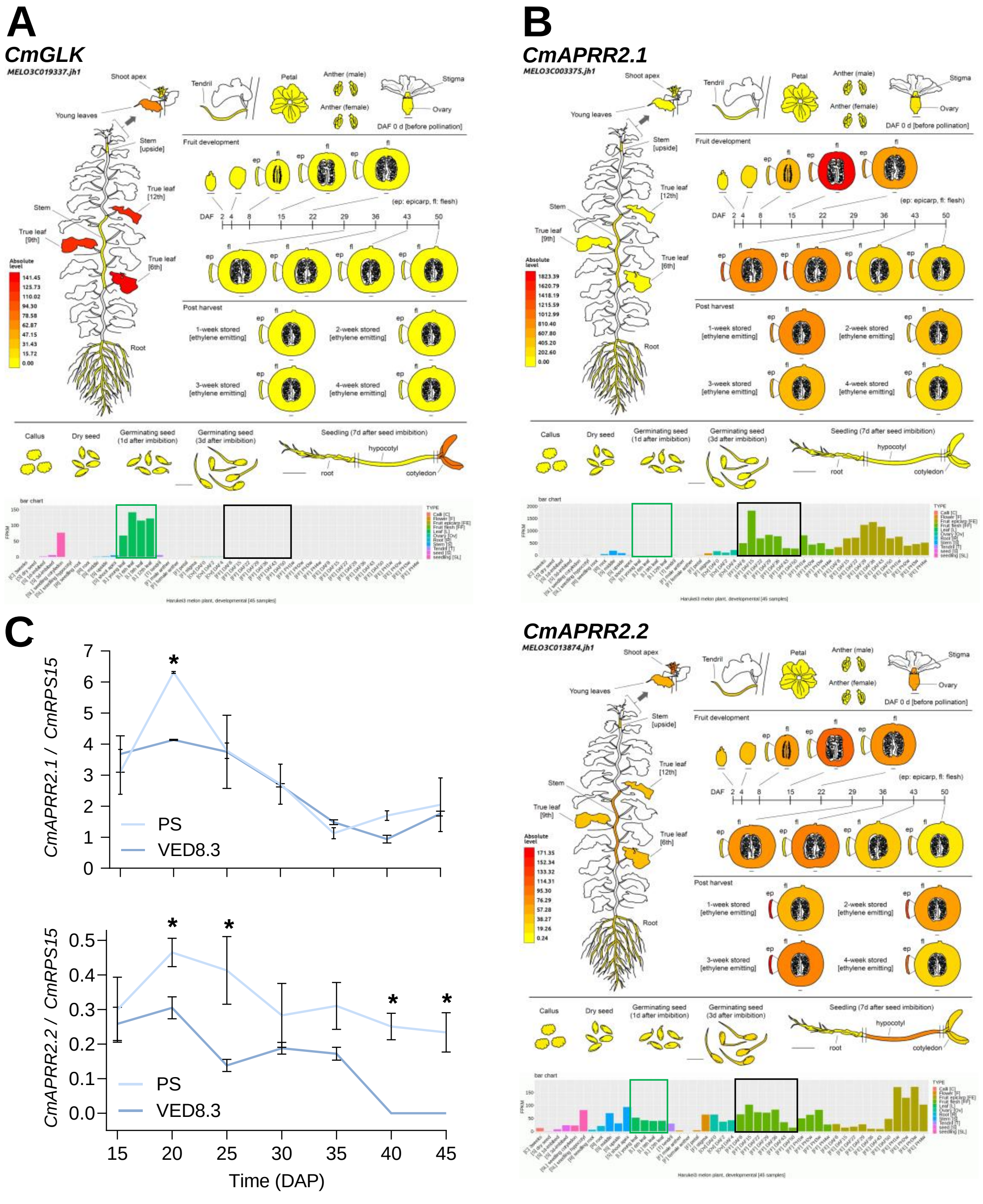

**Figure S13. Comparison of APRR2 homologs from different plants.** (A) Alignment of APRR2 proteins from *Arabidopsis thaliana* (*At*), *Benincasa hispida* (*Bh*, wax gourd), *Daucus carota* (*Dc*, carrot), *Capsicum annuum* (*Ca*, pepper), *Citrullus lanatus* (*Cl*, watermelon), *Cucumis melo* (*Cm*, melon), *Cucumis sativus* (*Cs*, cucumber), *Cucurbita pepo* (*Cp*, zucchini), *Cucurbita maxima* (*Cx*, pumpkin), *Lagenaria siceraria* (*Ls*, bottle gourd), *Momordica charantia* (*Mc*, bitter gourd), *Solanum lycopersicum* (*Sl*, tomato), and *Solanum melongena* (*Sm*, eggplant). Cm.1 corresponds to CmAPRR2.1 and Cm.2 corresponds to CmAPRR2.2. (B) Genetic distance among the proteins aligned in panel A.

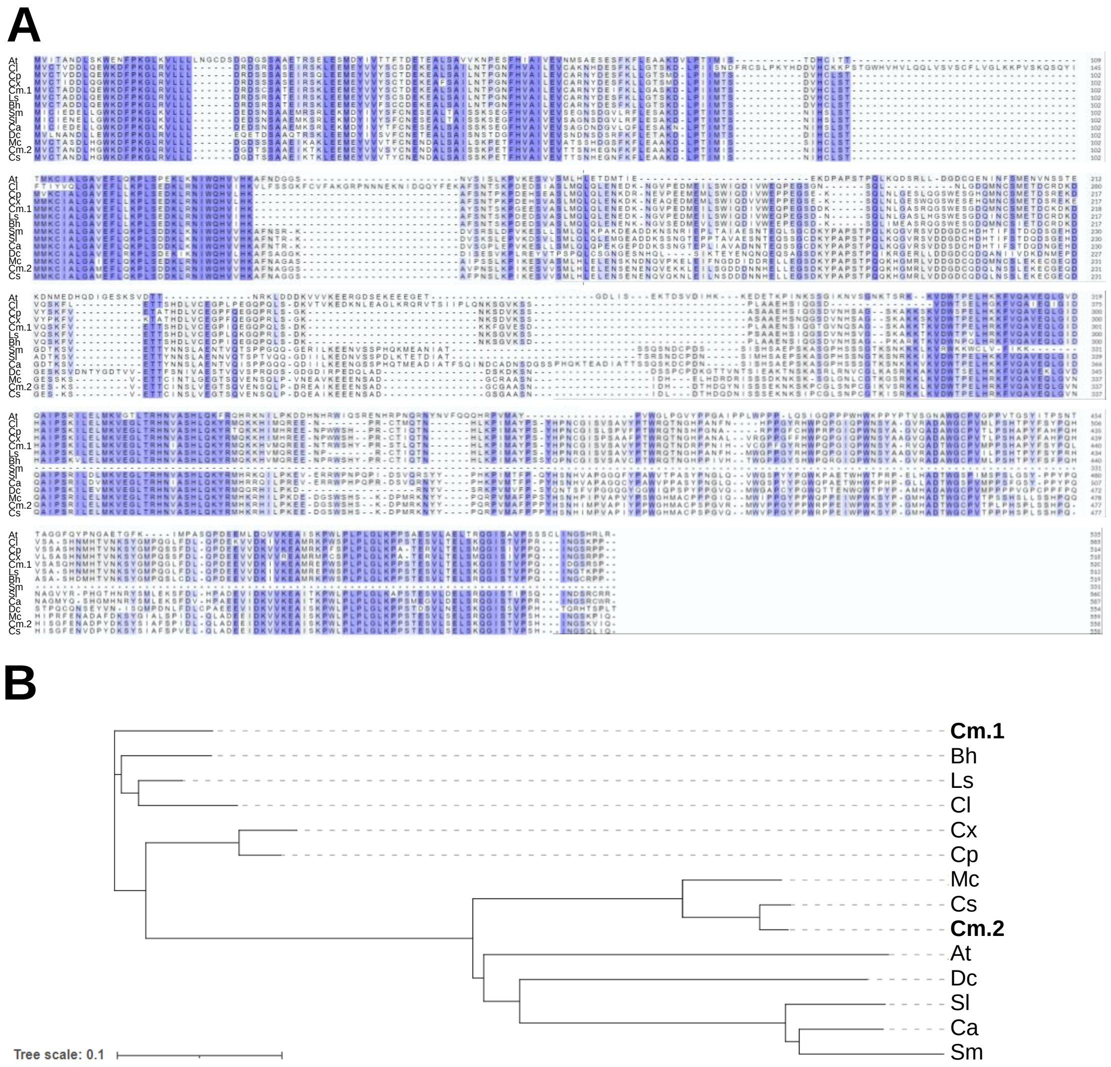

### SUPPLEMENTARY TABLES

#### Table S1. List of primers used for fine mapping.

| Primer name | Sequence | Length | Linked gene* |
|---|---|---|---|
| **chr08_31897002.A1** | GAAGGTGACCAAGTTCATGCTGCTAACCACTCGTGCGTC | 39 | |
| **chr08_31897002.A2** | GAAGGTCGGAGTCAACGGATTGCTGCTAACCACTCGTGCGTT | 42 | MELO3C003083 |
| **chr08_31897002.C1** | GCTGTTGTTACGCTTGATTCCATTGTAAT | 29 | |
| **chr08_31906121.A1** | GAAGGTGACCAAGTTCATGCTAATAGCATGACATGAAAAATAAACATGGATAA | 53 | |
| **chr08_31906121.A2** | GAAGGTCGGAGTCAACGGATTAATAGCATGACATGAAAAATAAACATGGATAT | 53 | MELO3C003086 |
| **chr08_31906121.C1** | CGTGTAGAAACTTGAAATTGTTCTCTGCTT | 30 | |
| **chr08_31910427.A1** | GAAGGTGACCAAGTTCATGCTGCAAGAATAATGGGAACCAGGATG | 45 | |
| **chr08_31910427.A2** | GAAGGTCGGAGTCAACGGATTGCAAGAATAATGGGAACCAGGATT | 45 | MELO3C003086 |
| **chr08_31910427.C1** | CCCTACTATGTTCTTGTATTTGAGAGTCAA | 30 | |
| **chr08_31968731.A1** | GAAGGTGACCAAGTTCATGCTGGCAGTCCATCATTCCAAGTGTTT | 45 | |
| **chr08_31968731.A2** | GAAGGTCGGAGTCAACGGATTGCAGTCCATCATTCCAAGTGTTC | 44 | MELO3C0092 |
| **chr08_31968731.C1** | GTGAACAGGGCAATCGCATTAAATATGAA | 29 | |
| **chr08_31977365.A1** | GAAGGTGACCAAGTTCATGCTGTCTTCGTATGTGAGTGTGTCATCA | 46 | |
| **chr08_31977365.A2** | GAAGGTCGGAGTCAACGGATTCTTCGTATGTGAGTGTGTCATCG | 44 | MELO3C0093 |
| **chr08_31977365.C1** | GACCACGACCATCTACAGTCTTGTA | 25 | |
| **chr08_31981113.A1** | GAAGGTGACCAAGTTCATGCTCGGGTCCGGTACGATGCAC | 40 | |
| **chr08_31981113.A2** | GAAGGTCGGAGTCAACGGATTTCGGGTCCGGTACGATGCAA | 41 | MELO3C0094 |
| **chr08_31981113.C1** | CAAAGAAGCGTACCGGAAACAGAGAT | 26 | |
| **chr08_31990033.A1** | GAAGGTGACCAAGTTCATGCTCGACCAACCTTCATCTACCCCA | 43 | |
| **chr08_31990033.A2** | GAAGGTCGGAGTCAACGGATTGACCAACCTTCATCTACCCCG | 42 | MELO3C0095 |
| **chr08_31990033.C1** | ATAAACCTTCCGGCGGCGGCAA | 22 | |
| **chr08_31995228.A1** | GAAGGTGACCAAGTTCATGCTAGTAAATAAGATAACTCCCAGCAATCC | 48 | |
| **chr08_31995228.A2** | GAAGGTCGGAGTCAACGGATTAGTAAATAAGATAACTCCCAGCAATCG | 48 | MELO3C0096 |
| **chr08_31995228.C1** | AGAAGAAGGGTCCGTGGAGAACAAA | 25 | |
| **chr08:32000845.A1** | GAAGGTGACCAAGTTCATGCTAGTTTTGAAATCTAACTTTTCAAACTCAAAAA | 53 | |
| **chr08:32000845.A2** | GAAGGTCGGAGTCAACGGATTAGTTTTGAAATCTAACTTTTCAAACTCAAAAG | 53 | MELO3C003096-MELO3C003097 |
| **chr08:32000845.C1** | AGCACATACTCCAAGAAAACAAGACCTAT | 29 | |
| **chr08_32003744.A1** | GAAGGTGACCAAGTTCATGCTAATTGTGTAATGAATCCTGAAACTAGAGAA | 51 | |
| **chr08_32003744.A2** | GAAGGTCGGAGTCAACGGATTGTGTAATGAATCCTGAAACTAGAGAG | 47 | MELO3C003097 |
| **chr08_32003744.C1** | CTGCTGTCTTCATCACCCAGATTCAT | 26 | |
| **chr08_32012692.A1** | GAAGGTGACCAAGTTCATGCTAAAGAGAAAAAGAAAATACCATGTGGATGA | 51 | |
| **chr08_32012692.A2** | GAAGGTCGGAGTCAACGGATTAAAGAGAAAAAGAAAATACCATGTGGATGT | 51 | MELO3C003098 |
| **chr08_32012692.C1** | AATCATTCCCACCACCATGGACGAT | 25 | |
| **chr08:32024368.A1** | GAAGGTGACCAAGTTCATGCTATATGTGCTGCTGCTAATATTGCA | 45 | |
| **chr08:32024368.A2** | GAAGGTCGGAGTCAACGGATTCTATATGTGCTGCTGCTAATATTGCT | 47 | MELO3C003099 |
| **chr08:32024368.C** | GCATGAAAACTCCTCCCTATCCCAA | 25 | |
| **chr08:32026558.A1** | GAAGGTGACCAAGTTCATGCTGATAATAATTACAAATCTAACCTAAATTATAA | 53 | |
| **chr08:32026558.A2** | GAAGGTCGGAGTCAACGGATTGATAATAATTACAAATCTAACCTAAATTATAT | 53 | MELO3C003099-MELO3C003100 |
| **chr08:32026558.C1** | GTAATTTATTACTCTCTTGCTTATACTAAT | 30 | |
| **chr08:32032539.A1** | GAAGGTGACCAAGTTCATGCTAATATCAATTCAATCTTCAATGTCAATGCC | 51 | |
| **chr08:32032539.A2** | GAAGGTCGGAGTCAACGGATTGAAATATCAATTCAATCTTCAATGTCAATGCT | 53 | MELO3C003100-MELO3C003101 |
| **chr08:32032539.C1** | GGTGTCGTTTTGACATTTCCTTCGGTT | 27 | |
| **chr08:32034658.A1** | GAAGGTGACCAAGTTCATGCTCAACATAATGTTGAAGTTTTACCTGCTG | 49 | |
| **chr08:32034658.A2** | GAAGGTCGGAGTCAACGGATTCCAACATAATGTTGAAGTTTTACCTGCTA | 50 | MELO3C003101 |
| **chr08:32034658.C** | GAAGGCGTACAGCAGATACCTTGTT | 25 | |
| **chr08:32048321.A1** | GAAGGTGACCAAGTTCATGCTAGAGATCGGAGAGGTATTGAAGATG | 46 | |
| **chr08:32048321.A2** | GAAGGTCGGAGTCAACGGATTCAGAGATCGGAGAGGTATTGAAGATA | 47 | MELO3C003104 |
| **chr08:32048321.C** | CTTCTCTCCCGATTCCCTTCCCAA | 24 | |
| **chr08:32054628.A1** | GAAGGTGACCAAGTTCATGCTACTGTTGCTACGGTTCCCACTAT | 44 | |
| **chr08:32054628.A2** | GAAGGTCGGAGTCAACGGATTCTGTTGCTACGGTTCCCACTAG | 43 | MELO3C003105 |
| **chr08:32054628.C** | AGAATCAAAAGCATCAGAAGCCAAGCTAA | 29 | |
| **chr08:32074736.A1** | GAAGGTGACCAAGTTCATGCTGATCATCTATCAAACGGACCATCCA | 46 | |
| **chr08:32074736.A2** | GAAGGTCGGAGTCAACGGATTATCATCTATCAAACGGACCATCCG | 45 | MELO3C003106 |
| **chr08:32074736.C** | GCCTTCTTTAGCTTCAATTGTTGTGCTTT | 29 | |
| **chr08:32114215.A1** | GAAGGTGACCAAGTTCATGCTACTAAGATTTTAAGTCAATTTGGCTTAATTTC | 53 | |
| **chr08:32114215.A2** | GAAGGTCGGAGTCAACGGATTGACTAAGATTTTAAGTCAATTTGGCTTAATTTT | 54 | MELO3C003108 |
| **chr08:32114215.C** | AACCACGTTTGGTCTTTGGTTCTTTGTTT | 29 | |
| **chr08:32118380.A1** | GAAGGTGACCAAGTTCATGCTTACCACTGATGAAGAGCTTAGGG | 44 | |
| **chr08:32118380.A2** | GAAGGTCGGAGTCAACGGATTACCACTGATGAAGAGCTTAGGC | 43 | MELO3C003109 |
| **chr08:32118380.C** | GTTCGTCTCAGCTCAACTCTCACAT | 25 | |
| **chr08:32122119.A1** | GAAGGTGACCAAGTTCATGCTTGCAGCTCTACTTGTATTCCATCG | 45 | |
| **chr08:32122119.A2** | GAAGGTCGGAGTCAACGGATTCTTGCAGCTCTACTTGTATTCCATCA | 47 | MELO3C003110 |
| **chr08:32122119.C** | TATATGCAAGCTCCAAGGACAGATTCAA | 28 | |

* when two genes are indicated, primers are linked to the intergenic region between them

#### Table S2. List of primers used for cloning and RT-qPCR.

| Primer # | Primer name | Sequence | Length | Use |
|---|---|---|---|---|
| 1 | attB1_98-gF | **GGGGACAAGTTTGTACAAAAAAGCAGGCT**ACCACCATGGACGATGACCC | 49 | Cloning |
| 2 | attB2_98-STOP-gR | **GGGGACCACTTTGTACAAGAAAGCTGGGT**CCGCTGTTCCATTCCACCTA | 49 | Cloning |
| 3 | attB2_98-FUS-gR | **GGGGACCACTTTGTACAAGAAAGCTGGGT**TCCGCCTCAGAGACGACGTC | 49 | Cloning |
| 4 | attB1-98-cF | **GGGGACAAGTTTGTACAAAAAAGCAGGCT**CCATGGACGATGACCCCTC | 48 | Cloning |
| 5 | attB1_98VED-cF | **GGGGACAAGTTTGTACAAAAAAGCAGGCT**TCATGGTGCACCACCTGATCG | 50 | Cloning |
| 6 | attB1-APRR2.1 | **GGGGACAAGTTTGTACAAAAAAGCAGGCT**TCATGGTTTGCACTGCCGACG | 50 | Cloning |
| 7 | attB2-APRR2.1 | **GGGGACCACTTTGTACAAGAAAGCTGGGT**TGGGTGATCTGGAGCCGTCG | 49 | Cloning |
| 8 | attB1-APRR2.2 | **GGGGACAAGTTTGTACAAAAAAGCAGGCT**TCATGGTTTGCACTGCTAATG | 50 | Cloning |
| 9 | attB2-APRR2.2 | **GGGGACCACTTTGTACAAGAAAGCTGGGT**TTTGGATTACTTTAGAGCCA | 49 | Cloning |
| 10 | CmRPGE1_qF | TCTGCTGGGTTCGAGGGATA | 20 | RT-qPCR |
| 11 | CmRPGE1_qR | TCCCACCTAGCCTACGACAC | 20 | RT-qPCR |
| 12 | CmAPRR2.1_qF | ACCAGATGAGGAGGTGGTTG | 20 | RT-qPCR |
| 13 | CmAPRR2.1_qR | GTCGATTTGAGGAGGGACGG | 20 | RT-qPCR |
| 14 | CmAPRR2.2_qF | TGAAAACGCCGATCCATACG | 20 | RT-qPCR |
| 15 | CmAPRR2.2_qR | CCATGGCTTGCTGATTGCC | 19 | RT-qPCR |
| 16 | CmRPS15_NqF | GAAGCTGCGTAAAGCGAAAC | 20 | RT-qPCR |
| 17 | CmRPS15_NqR | GGTCTTTCCATTGTAAACTCCAA | 23 | RT-qPCR |
| 18 | NbORE1-qF | GGGCAAAACATGCAGCAGAA | 20 | RT-qPCR |
| 19 | NbORE1-qR | AGTCCAGAGGCAATCAAGATCC | 22 | RT-qPCR |
| 20 | NbACT-NqF | TAAGGTTGTTGCACCACCAG | 20 | RT-qPCR |
| 21 | NbACT-NqR | ACATCTGCTGGAATGTGCTG | 20 | RT-qPCR |

#### Table S3. Constructs generated in this work.

| F* | R* | Template | Plasmid | Insert | Use |
|---|---|---|---|---|---|
| 1 | 2 | gDNA | pGWB605 | CmRPGE1$^{PS}$-STOP | Activity assays in *N. benthamiana* (Fig. 3), transgenic lines in A. thaliana (Fig. 4) |
| 1 | 2 | gDNA | pGWB605 | CmRPGE1$^{VED}$-STOP | Activity assays in *N. benthamiana* (Fig. 3), transgenic lines in A. thaliana (Fig. 4) |
| 1 | 3 | gDNA | pGWB605 | CmRPGE1$^{PS}$-GFP | Activity assays and subcellular localization in *N. benthamiana* (Fig. 3) |
| 1 | 3 | gDNA | pGWB605 | CmRPGE1$^{VED}$-GFP | Activity assays and subcellular localization in *N. benthamiana* (Fig. 3) |
| 4 | 3 | gDNA | pGWB454 | CmRPGE1$^{PS}$-RFP | Subcellular localization in *N. benthamiana* (Fig. 3) |
| 5 | 3 | gDNA | pGWB454 | CmRPGE1$^{VED}$-RFP | Subcellular localization in *N. benthamiana* (Fig. 3) |
| 4 | 3 | cDNA | pGWB605 | CmRPGE1$^{PS}$-GFP | Subcellular localization in *N. benthamiana* (Fig. 5) |
| 5 | 3 | cDNA | pGWB605 | CmRPGE1$^{VED}$-GFP | Subcellular localization in *N. benthamiana* (Fig. 5) |
| 6 | 7 | cDNA | pGWB454 | CmAPRR2.1-RFP | Subcellular localization in *N. benthamiana* (Fig. 5) |
| 8 | 9 | cDNA | pGWB454 | CmAPRR2.2-RFP | Subcellular localization in *N. benthamiana* (Fig. 5) |
| 4 | 3 | cDNA | pGWB420 | CmRPGE1$^{PS}$-myc | Expression in *N. benthamiana* and inmunoprecipitation (Fig. 6) |
| 5 | 3 | cDNA | pGWB420 | CmRPGE1$^{VED}$-myc | Expression in *N. benthamiana* and inmunoprecipitation (Fig. 6) |
| 6 | 7 | cDNA | pGWB414 | CmAPRR2.1-HA | Expression in *N. benthamiana* and inmunoprecipitation (Fig. 6) |
| 8 | 9 | cDNA | pGWB414 | CmAPRR2.2-HA | Expression in *N. benthamiana* and inmunoprecipitation (Fig. 6) |
| 4 | 3 | cDNA | YFC43-GW | (C)YFP-CmRPGE1$^{PS}$ | BiFC in *N. benthamiana* (Fig. 5) |
| 5 | 3 | cDNA | YFC43-GW | (C)YFP-CmRPGE1$^{VED}$ | BiFC in *N. benthamiana* (Fig. 5) |
| 6 | 7 | cDNA | YFN43-GW | (N)YFP-CmAPRR2.1 | BiFC in *N. benthamiana* (Fig. 5) |
| 8 | 9 | cDNA | YFN43-GW | (N)YFP-CmAPRR2.2 | BiFC in *N. benthamiana* (Fig. 5) |

* Numbers correspond to primers in Table S2
